# Chromosome painting in cultivated banana and their wild relatives (*Musa* spp.) reveals differences in chromosome structure

**DOI:** 10.1101/2020.08.01.232207

**Authors:** D Šimoníková, A Němečková, J Čížková, A Brown, R Swennen, J Doležel, E Hřibová

**Author notes:** Corresponding author: Eva Hřibová, Institute of Experimental Botany, Centre of the Region Hana for Biotechnological and Agricultural Research, Šlechtitelů 31, CZ-77900 Olomouc, Czech Republic.

## Abstract

Edible banana cultivars are diploid, triploid or tetraploid hybrids which originated by natural cross hybridization between subspecies of diploid *Musa acuminata*, or between *M. acuminata* and diploid *M. balbisiana*. Participation of two other wild diploid species *M. schizocarpa* and *M. textilis* was also indicated by molecular studies. Fusion of gametes with structurally different chromosome sets may give rise to progenies with structural chromosome heterozygosity and reduced fertility due to aberrant chromosome pairing and unbalanced chromosome segregation. Only a few translocations have been classified on the genomic level so far and a comprehensive molecular cytogenetic characterization of cultivars and species of the family *Musaceae* is still lacking. FISH with chromosome-arm specific oligo painting probes was used for comparative karyotype analysis in a set of wild *Musa* species and edible banana clones. The results revealed large differences in chromosome structure discriminating individual accessions. These results permitted identification of putative progenitors of cultivated clones and clarified genomic constitution and evolution of aneuploid banana clones, which seem to be common among the polyploid banana accessions. New insights into the chromosome organization and structural chromosome changes will be a valuable asset in breeding programs, particularly in selection of appropriate parents for cross hybridization.

**Highlight:** Oligo painting FISH revealed chromosomal translocations in subspecies of *Musa acuminata* (A genome), their intra-specific hybrids as well as in *M. balbisiana* (B genome) and in interspecific hybrid clones originating from cross hybridization between *M. acuminata* and *M. balbisiana*

## Introduction

Banana represents one of the major staple foods and is one of the most important cash crops with the estimated value of $25 billion for the banana industry. An annual global production of bananas reached 114 million tons in 2017 (FAOSTAT, 2017) with about 26 million tons exported in 2019 (International Trade Statistics). Two types of bananas are known - sweet bananas, serving as a food supplement, and cooking bananas, which are characteristic by starchier fruits (Price, 1995). Edible banana cultivars are vegetatively propagated diploid, triploid and tetraploid hybrids which originated after natural cross hybridization between wild diploids *Musa acuminata* (2n=2x=22, AA genome) and *M. balbisiana* (2n=2x=22, BB genome) and their hybrid progenies. To some extent, also other *Musa* species such as *M. schizocarpa* (2n=22=22, SS genome) and *M. textilis* (2n=2x=20, TT genome) contributed to the origin of some edible banana clones (Carreel *et al*., 1994, Čížková *et al*., 2013, Němečková *et al*., 2018).

Based on morphology and geographical distribution, *M. acuminata* has been divided into nine subspecies (*banksii, burmannica, burmannicoides, errans, malaccensis, microcarpa, siamea, truncata* and *zebrina*) and three varieties (chinensis, sumatrana, tomentosa) (Perrier *et al*., 2011, Martin *et al*., 2017, *WCSP, 2018*). It has been estimated that least four subspecies of *M. acuminata* contributed to the origin of cultivated bananas (Perrier *et al*., 2011, Rouard *et al*., 2018). Out of them, *M. acuminata* ssp. *banksii*, with the original center of diversity in New Guinea played a major role in this process (Sharrock, 1990, Perrier *et al*., 2009, Němečková *et al*., 2018). Other subspecies were *M. acuminata* ssp. *burmannica* with the center of diversity in Myanmar (Cheesman, 1948), ssp. *malaccensis* with the origin in Malay peninsula (De Langhe *et al*., 2009, Perrier *et al*., 2011) and ssp. *zebrina* which originated in Indonesia (Rouard *et al*., 2018).

It is believed that human migration together with a different geography of the present archipelago in South-East Asia during glacial period, when drop of sea level resulted in interconnection of current islands in South-East Asia into one land mass (Sand, 1989, Denham, 2004, Denham, 2010, Kagy *et al*., 2016), brought different *M. acuminata* subspecies to a close vicinity, enabling cross hybridization and giving rise to diploid intra-specific hybrids that were subjected to human selection and propagation (Perrie*r et al*., 2011, Martin *et al*., 2017). Fusion of unreduced gametes produced by diploid edible and partially sterile cultivars with normal haploid gametes from fertile diploid (Simmonds, 1962, Carreel *et al*., 1994, Raboin *et al*., 2005) would give rise to triploids. The number one export dessert banana type Cavendish as well as other important dessert bananas, such as Gros Michel and Pome types, originated according to this scenario by hybridization of a diploid representative of sub-group ‘Mchare’ (originally named ‘Mlali’) (AA genome; *zebrina / microcarpa* and *banksii* ascendance) which served as a donor of an unreduced diploid gamete with haploid gamete of ‘Khai’ (*malaccensis* ascendance) (Perrier *et al*., 2009, Perrier *et al*., 2019, Martin *et al*., 2020).

Another group of edible triploid bananas, clones with AAB (so called plantains) or ABB constitution, cover nearly 40 % of global banana production, whereas plantains stand for 18 % of total banana production (Baurens *et al*., 2019). These interspecific triploid cultivars originated after fusion of an unreduced gamete from interspecific AB hybrid with haploid gamete from diploid *M. acuminata* ssp. *banksii* or *M. balbisiana* (Perrier *et al*., 2011, Baurens *et al*., 2019). Their evolution most probably involved several backcrosses (De Langhe *et al*., 2010). Two important AAB subgroups of starchy bananas evolved in two centers of diversity, the African Plantains and the Pacific (Maia Maoli / Popoulu) Plantains (De Langhe *et al*., 2009).

East African Highland bananas (EAHB) represent an important starchy type of banana for over 80 million people living in the Great Lakes region of East Africa, which is considered as a secondary center of banana diversity (Cooper *et al*., 2001, Tugume *et al*., 2003). EAHBs are a sub-group of triploid bananas with AAA constitution, which arose from hybridization between diploid *M. acuminata* ssp. *banksii* and *M. acuminata* ssp. *zebrina*. However, also *M. schizocarpa* contributed to the formation of these hybrids (Němečková *et al*., 2018). Interestingly, different EAHB varieties have relatively low genetic diversity, on the contrary to morphological variation which occurred most probably due to the accumulation of somatic mutations and epigenetic changes (Perrier *et al*., 2009, Perrier *et al*., 2011, Kitavi *et al*., 2016, Christelová *et al*., 2017, Němečková *et al*., 2018).

Cultivated clones originating from inter-subspecific and inter-specific hybridization and with a contribution of unreduced gametes in case of triploid clones have reduced or zero production of fertile gametes. This is a consequence of aberrant chromosome pairing during meiosis due to structural chromosome heterozygosity and/or odd ploidy levels. Reduced fertility greatly hampers the efforts to breed improved cultivars (Burke and Arnold, 2001, Martin *et al*., 2017, Baurens *et al*., 2019, Batte *et al*., 2019), which are needed to satisfy the increasing demand for dessert and starchy bananas under the conditions of climate change and increasing pressure of pests and diseases (Christelová *et al*., 2017).

Traditional breeding strategies for triploid banana cultivars involve development of tetraploids (4x) from 3x × 2x crosses, followed by production of secondary triploid hybrids (3x) from 4x × 2x crosses (Bakry and Horry, 1992, Tomepke *et al*., 2004, Ortiz, 2013, Nyine *et al*., 2017). Similarly, diploids play important role also in breeding strategy of diploid cultivars, 4x × 2x crosses are used to create improved cultivars (Ortiz, 2013). It is thus necessary to identify cultivars that produce seeds under specific condition, followed by breeding for target traits and re-establishing seed-sterile end product. Low seed yield after pollination (e.g. 4 seeds per Matooke (genome AAA) bunch and only 1.6 seeds per Mchare (genome AA) bunch followed by embryo rescue, with very low germination rate (∼ 2%) illustrates the serious bottleneck for the breeding processes (Brown and Swennen, unpublished).

Since banana breeding programs use diploids as the principle vehicle for introducing genetic variability (e.g. Amorim *et al*., 2011, Amorim *et al*., 2013, Tenkouano *et al*., 2003), the knowledge of their genome structure at chromosomal level is critical to reveal possible causes of reduced fertility, and presence of non-recombining haplotype blocks, provide data to identify parents of cultivated clones, which originated spontaneously without a direct human intervention, and to select parents for improvement programs. In order to provide insights into the genome structure at chromosome level, we employed FISH with chromosome-arm specific oligo painting probes in a set of wild *Musa* species and edible banana clones potentially useful in banana improvement. Chromosome painting in twenty representatives of the Eumusa section of genus *Musa*, which included subspecies of *M. acuminata, M. balbisiana* and their inter-subspecific and inter-specific hybrids, revealed chromosomal rearrangements discriminating subspecies of *M. acuminata* and structural chromosome heterozygosity of cultivated clones. Identification of chromosome translocations pointed to particular *Musa* subspecies as putative parents of cultivated clones and provided an independent support for hypotheses on their origin.

## Materials and Methods

### Plant material and diversity tree construction

Representatives of twenty species and clones from the section Eumusa of genus *Musa* were obtained as *in vitro* rooted plants from the International *Musa* Transit Centre (ITC, Bioversity International, Leuven, Belgium). *In vitro* plants were transferred to soil and kept in a heated greenhouse. Table 1 lists the accessions used in this work. Genetic diversity analysis of banana accessions used in the study was performed using SSR data according to Christelová *et al*. (2017). Dendrogram was constructed based on the results of UPGMA analysis implemented in DARwin software v6.0.021 (Perrier and Jacquemoud-Collet, 2006) and visualized in FigTree v1.4.0 (http://tree.bio.ed.ac.uk/software/figtree/).

**Table 1:** List of *Musa* accessions analyzed in this work and their genomic constitution

| Species | Subspecies / Subgroup | Accession name | ITC code<sup>a</sup> | Genomic constitution | Chromosome number (2n) |
| --- | --- | --- | --- | --- | --- |
| *M. acuminata* | *banksii* | Banksii | 0806 | AA | 22 |
|  | *microcarpa* | Borneo | 0253 | AA | 22 |
|  | *zebrina* | Maia Oa | 0728 | AA | 22 |
|  | *burmannica* | Tavoy | 0072 | AA | 22 |
|  | *burmannicoides* | Calcutta 4 | 0249 | AA | 22 |
|  | *siamea* | Pa Rayong | 0672 | AA | 22 |
| ***M. balbisiana*** | - | Pisang Klutuk Wulung | - | BB | 22 |
| **Cultivars** | unknown | Pisang Lilin | - | AA | 22 |
|  | Mchare | Huti White |  | AA | 22 |
|  | Mchare | Huti (Shumba nyeelu) | 1452 | AA | 22 |
|  | Mchare | Ndyali | 1552 | AA | 22 |
|  | Cavendish | Poyo | 1482 | AAA | 33 |
|  | Gros Michel | Gros Michel | 0484 | AAA | 33 |
|  | Mutika/Lujugira | Imbogo | 0168 | AAA | 32* |
|  | Mutika/Lujugira | Kagera | 0141 | AAA | 33 |
|  | Plantain (Horn) | 3 Hands Planty | 1132 | AAB | 33 |
|  | Plantain (French) | Amou | 0963 | AAB | 32* |
|  | Plantain (French) | Obino l'Ewai | 0109 | AAB | 33 |
|  | Pelipita | Pelipita | 0472 | ABB | 33 |
|  | Saba | Saba sa Hapon | 1777 | ABB | 33 |

<sup>a</sup>Code assigned by the International Transit Centre (ITC, Leuven, Belgium).
\*Aneuploidy observed in our study.

### Preparation of oligo painting probes and mitotic metaphase chromosome spreads

Chromosome-arm specific painting probes were prepared as described by Šimoníková *et al*. (2019). Briefly, sets of 20,000 oligomers (45-nt) covering individual chromosome arms were synthesized as immortal libraries by Arbor Biosciences (Ann Arbor, Michigan, USA) and then labeled directly by CY5 fluorochrome, or by digoxigenin or biotin according to Han *et al*. (2015). N.B.: In the reference genome assembly of *M. acuminata* DH Pahang (D’Hont *et al*., 2012), pseudomolecules 1, 6, 7 are oriented inversely to the traditional way karyotypes are presented, where the short arms are on the top (Suppl. Fig. S3, see also Šimoníková *et al*., 2019).

Actively growing root tips were collected and pre-treated in 0.05% (w/v) 8-hydroxyquinoline for three hours at room temperature, fixed in 3:1 ethanol:acetic acid fixative overnight at −20°C and stored in 70% ethanol at −20°C. After washing in 75-mM KCl and 7.5-mM EDTA (pH 4), root tip segments were digested in a mixture of 2% (w/v) cellulase and 2% (w/v) pectinase in 75-mM KCl and 7.5-mM EDTA (pH 4) for 90 min at 30°C. The suspension of protoplasts thus obtained was filtered through a 150-μm nylon mesh, pelleted and washed in 70% ethanol. For further use, the protoplast suspension was stored in 70% ethanol at −20°C. Mitotic metaphase chromosome spreads were prepared by a dropping method from protoplast suspension according to Doležel *et al*. (1998), the slides were postfixed in 4% (v/v) formaldehyde solution in 2 × SSC solution, air dried and used for FISH.

### Fluorescence *in situ* hybridization and image analysis

Fluorescence *in situ* hybridization and image analysis were performed according to Šimoníková *et al*. (2019). Hybridization mixture containing 50% (v/v) formamide, 10% (w/v) dextran sulfate in 2 × SSC and 10 ng/µl of labeled probes was added onto slide and denatured for 3 min at 80°C. Hybridization was carried out in a humid chamber overnight at 37°C. The sites of hybridization of digoxigenin- and biotin-labeled probes were detected using anti-digoxigenin-FITC (Roche Applied Science, Penzberg, Germany) and streptavidin-Cy3 (ThermoFisher Scientific/Invitrogen, Carlsbad, CA, USA), respectively. Chromosomes were counterstained with DAPI and mounted in Vectashield Antifade Mounting Medium (Vector Laboratories, Burlingame, CA, USA). The slides were examined with Axio Imager Z.2 Zeiss microscope (Zeiss, Oberkochen, Germany) equipped with Cool Cube 1 camera (Metasystems, Altlussheim, Germany) and appropriate optical filters. The capture of fluorescence signals, merging the layers and measurement of chromosome length were performed with ISIS software 5.4.7 (Metasystems). The final image adjustment and creation of idiograms were done in Adobe Photoshop CS5.

## Results

### Karyotype of *M. acuminata* and *M. balbisiana*

The first part of the study focused on comparative karyotype analysis in six subspecies of *M. acuminata* and in *M. balbisiana*. Oligo painting FISH in structurally homozygous *M. acuminata* ssp. *banksii* ‘Banksii’ and *M. acuminata* ssp. *microcarpa* ‘Borneo’ did not reveal detectable chromosome translocations (Fig. 1A) as compared to the reference banana genome of *M. acuminata* ssp. *malaccensis* ‘DH Pahang’ (Šimoníková *et al*., 2019). Thus, the three subspecies share the same overall organization of their chromosome sets. On the other hand, different chromosome translocations, which were found to be subspecies-specific were identified in the remaining four subspecies of *M. acuminata* (ssp. *zebrina*, ssp. *burmannica*, ssp. *burmannicoides*, ssp. *siamea*).

**Figure 1:**
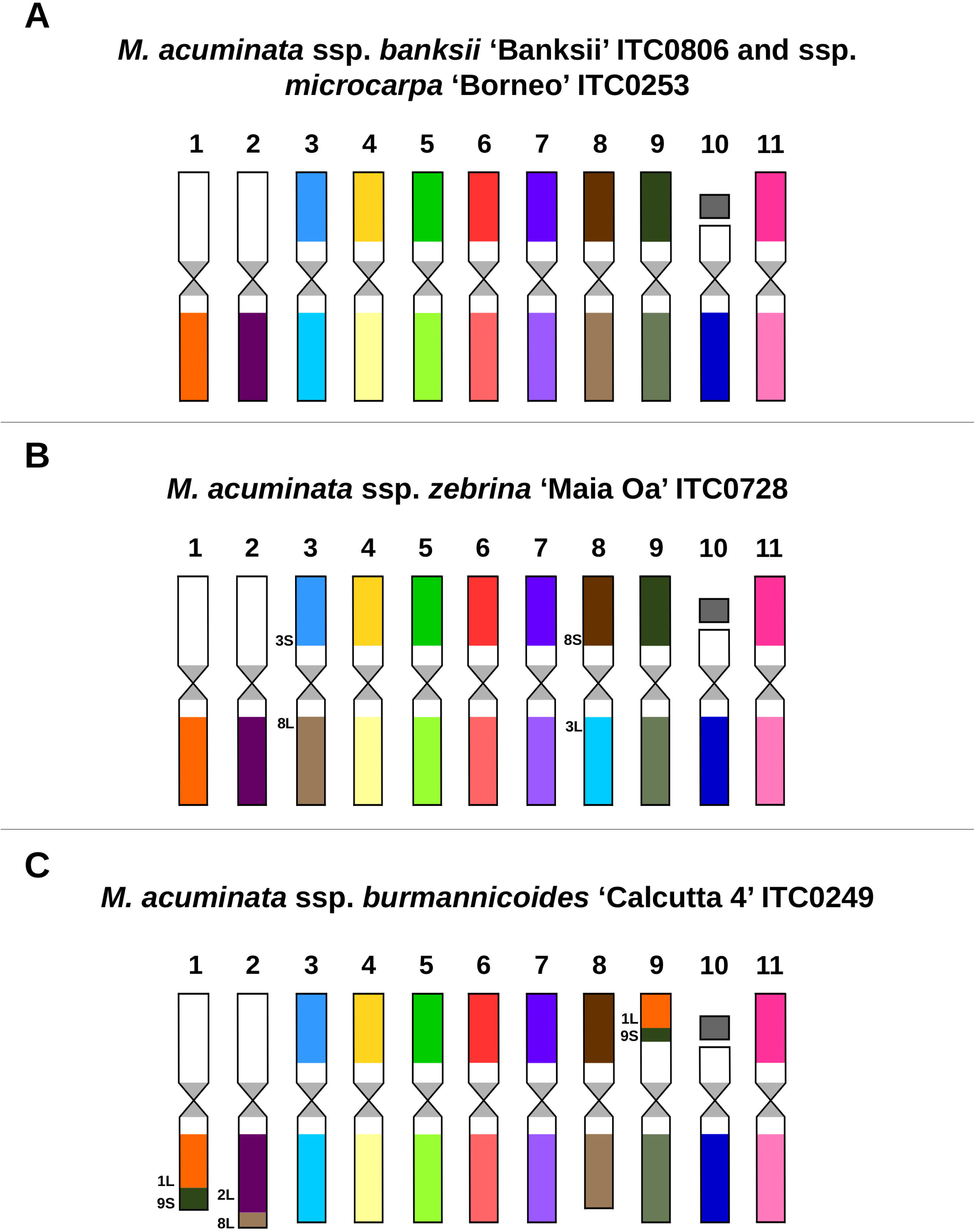
Idiograms of four structurally homozygous diploid (2n=2x=22) subspecies of *M. acuminata*: (A) *M. acuminata* ssp. *banksii* ‘Banksii’ and ssp. *microcarpa* ‘Borneo’; (B) *M. acuminata* ssp. *zebrina* ‘Maia Oa’; (C) *M. acuminata* ssp. *burmannicoides* ‘Calcutta 4’. Chromosomes are oriented with their short arms of at the top. Chromosome paints were not used for short arms of chromosomes 1, 2 and 10.

In *M. acuminata* ssp. *zebrina*, a reciprocal translocation between the short arm of chromosome 3 and the long arm of chromosome 8 was identified (Figs. 1B, 2A). Three phylogenetically closely related subspecies *M. acuminata* ssp. *burmannica, M. acuminata* ssp. *burmannicoides* and *M. acuminata* ssp. *siamea* shared two translocations (Figs. 1C, 3A, 3B, 3E, 3F; Suppl. Figs. S1A, S1B). The first of these involved a transfer of a segment of the long arm of chromosome 8 to the long arm of chromosome 2; the second was a reciprocal translocation involving a large segment of the short arm of chromosome 9 and the long arm of chromosome 1. In addition to the translocations shared by representatives of the three subspecies, chromosome painting revealed additional subspecies-specific translocations.

**Figure 2:**
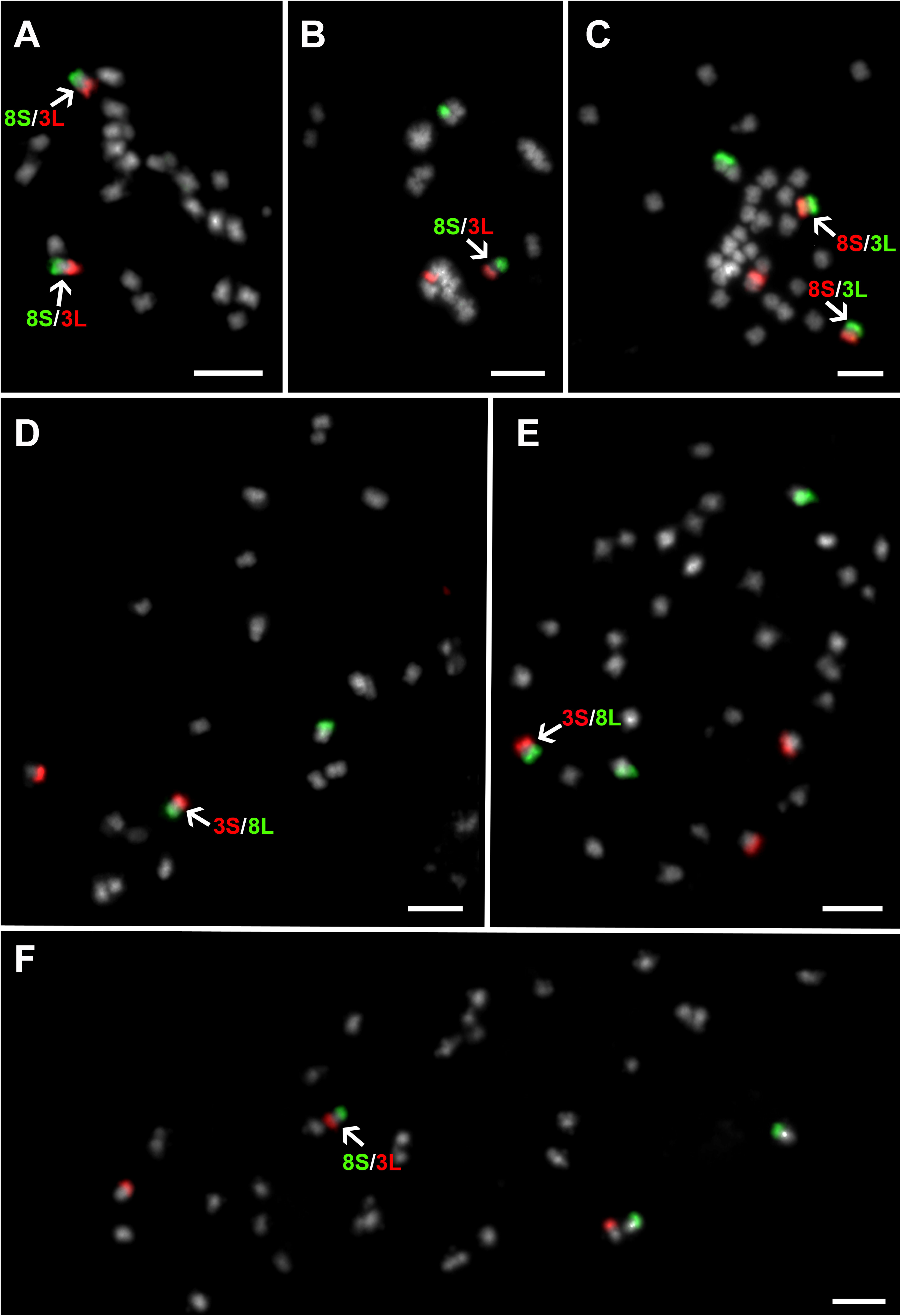
Examples of oligo painting FISH on mitotic metaphase plates of six *Musa* accessions: (A) ‘Maia Oa’ (2n=2x=22, AA genome), probes for long arm of chromosome 3 and short arm of chromosome 8 were labeled in red and green, respectively; (B) ‘Tavoy’ (2n=2x=22, AA genome), probes for long arm of chromosome 3 and short arm of chromosome 8 were labeled in red and green, respectively; (C) ‘Imbogo’ (2n=3x - 1= 32, AAA genome): probes for long arm of chromosome 3 and short arm of chromosome 8 were labeled in greed and red, respectively; (D) ‘Huti (Shumba nyeelu)’ (2n=2x=22, AA genome), probes for short arm of chromosome 3 and long arm of chromosome 8 were labeled in red and green, respectively; (E) ‘Gros Michel’ (2n=3x=33, AAA genome), probes for short arm of chromosome 3 and long arm of chromosome 8 were labeled in red and green, respectively; (F) ‘Poyo’ (2n=2x=33, AAA genome), probes for long arm of chromosome 3 and short arm of chromosome 8 were labeled in red and green, respectively. Chromosomes were counterstained with DAPI (light grey pseudocolor). Arrows point translocation chromosomes. Bars = 5 µm.

**Figure 3:**
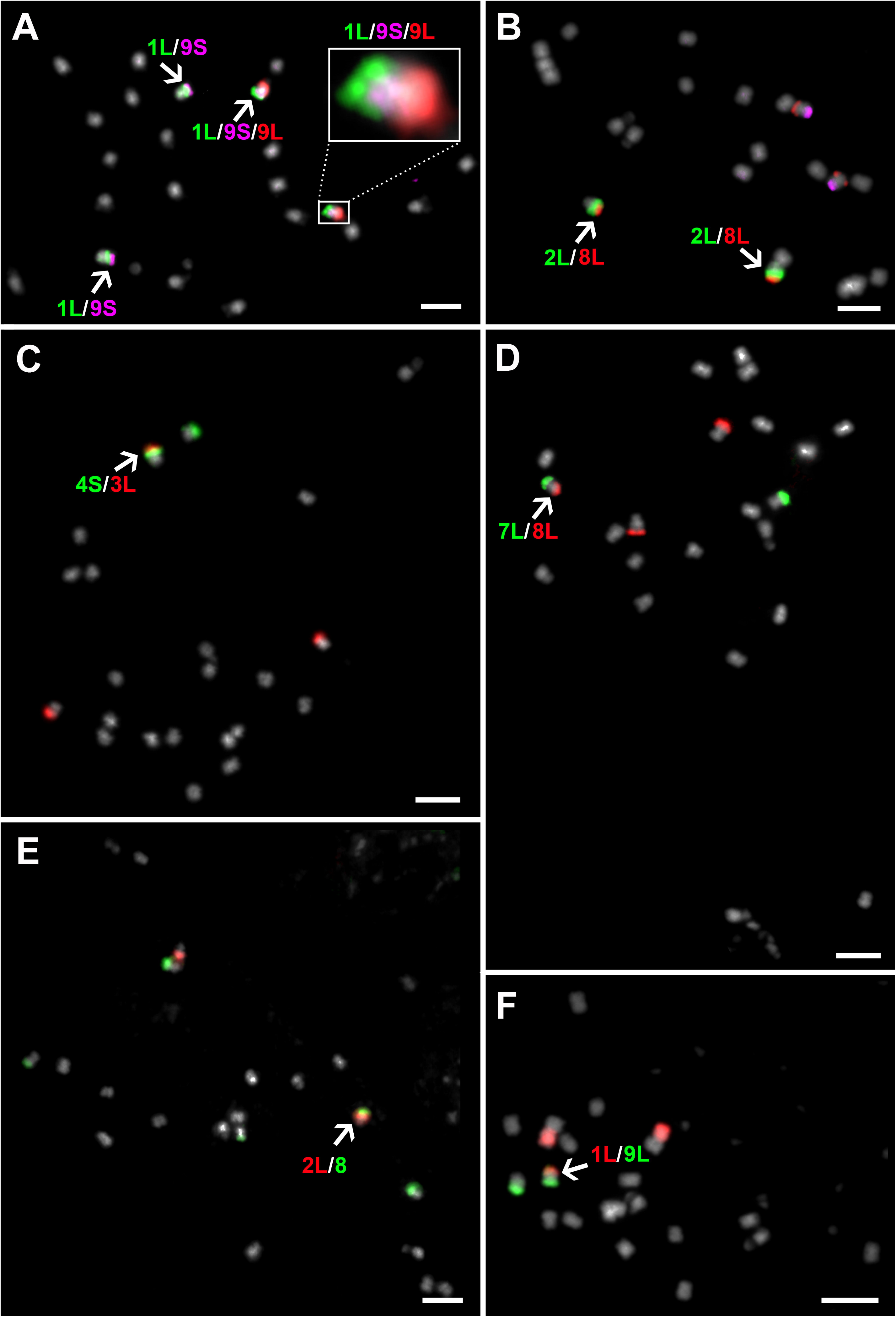
Examples of oligo painting FISH on mitotic metaphase plates of six *Musa* accessions: (A) ‘Calcutta 4’ (2n=2x=22, AA genome), probes for long arm of chromosome 1, long arm of chromosome 9 and short arm of chromosome 9 were labeled in green, red and purple, respectively; (B) ‘Calcutta 4’ (2n=2x=22, AA genome), probes for long arm of chromosome 2, long arm of chromosome 8 and short arm of chromosome 8 were labeled in green, red and purple, respectively; (C) ‘Pa Rayong’ (2n=2x=22, AA genome), probes for long arm of chromosome 3 and short arm of chromosome 4 were labeled in red and green, respectively; (D) ‘Tavoy’ (2n=2x=22, AA genome), probes for long arm of chromosome 7 and long arm of chromosome 8 were labeled in green and red, respectively; (E) ‘Tavoy’ (2n=2x=22, AA genome), probes for long arm of chromosome 2 and chromosome 8 were labeled in red and green, respectively; (F) ‘Tavoy’ (2n=2x=22, AA genome), probes for long arm of chromosome 1 and long arm of chromosome 9 were labeled in red and green, respectively. Chromosomes were counterstained with DAPI (light grey pseudocolor). Arrows point translocation chromosomes. Bars = 5 µm.

In *M. acuminata* ssp. *siamea* ‘Pa Rayong’ translocation of a small segment of the long arm of chromosome 3 to the short arm of chromosome 4 was detected. Importantly, this translocation was visible only on one of the homologs, indicating structural chromosome heterozygosity and a hybrid origin (Fig. 3C; Suppl. Fig. S1A). Subspecies-specific translocations involving only one of the homologs were also found in *M. acuminata* ssp. *burmannica* ‘Tavoy’. They included a reciprocal translocation between chromosomes 3 and 8, which was also detected in *M. acuminata* ssp. *zebrina*, and a Robertsonian translocation between chromosomes 7 and 8, which gave rise to a chromosome comprising long arms of chromosomes 7 and 8 and a chromosome made of short arms of chromosomes 7 and 8 (Figs. 2B, 3D; Suppl. Fig. S1B).

In *M. balbisiana* ‘Pisang Klutuk Wulung’ translocation of a small segment of the long arm of chromosome 3 to the long arm of chromosome 1 was observed (Suppl. Fig. S1C), similar to the translocation identified in our previous work (Šimoníková *et al*., 2019) in *M. balbisiana* ‘Tani’. In agreement with a low level of genetic diversity of *M. balbisiana* (De Langhe *et al*., 2015, Kagy *et al*., 2016), no detectable differences in karyotypes were found between both accessions of the species.

### Karyotype structure of edible clones which originated as intra-specific hybrids

Chromosome painting in diploid cooking banana cultivars belonging to the Mchare group with AA genome, confirmed their hybrid origin. All three accessions analyzed in this work comprised two reciprocal translocations, with each of them observed only in one chromosome set. The first translocation involved short segments of long arms of chromosomes 4 and 1 (Suppl. Fig. S1E), while the second one involved short arms of chromosomes 3 and long arm of chromosome 8. Both translocations were observed in heterozygous state in all three analyzed Mchare banana representatives (Fig. 2D; Suppl. Fig. S1E). Reciprocal translocation involving short segments of long arms of chromosomes 4 and 1 was also identified in diploid cultivar ‘Pisang Lilin’ with AA genome (Fig. 4C; Suppl. Fig. S1D). This clone is used in breeding programs as a donor of useful agronomical characteristics, such as female fertility or resistance to yellow Sigatoka (do Amaral *et al*., 2015). Also this accession was heterozygous for the translocation, indicating a hybrid origin (Suppl. Fig. S1D).

**Figure 4.**
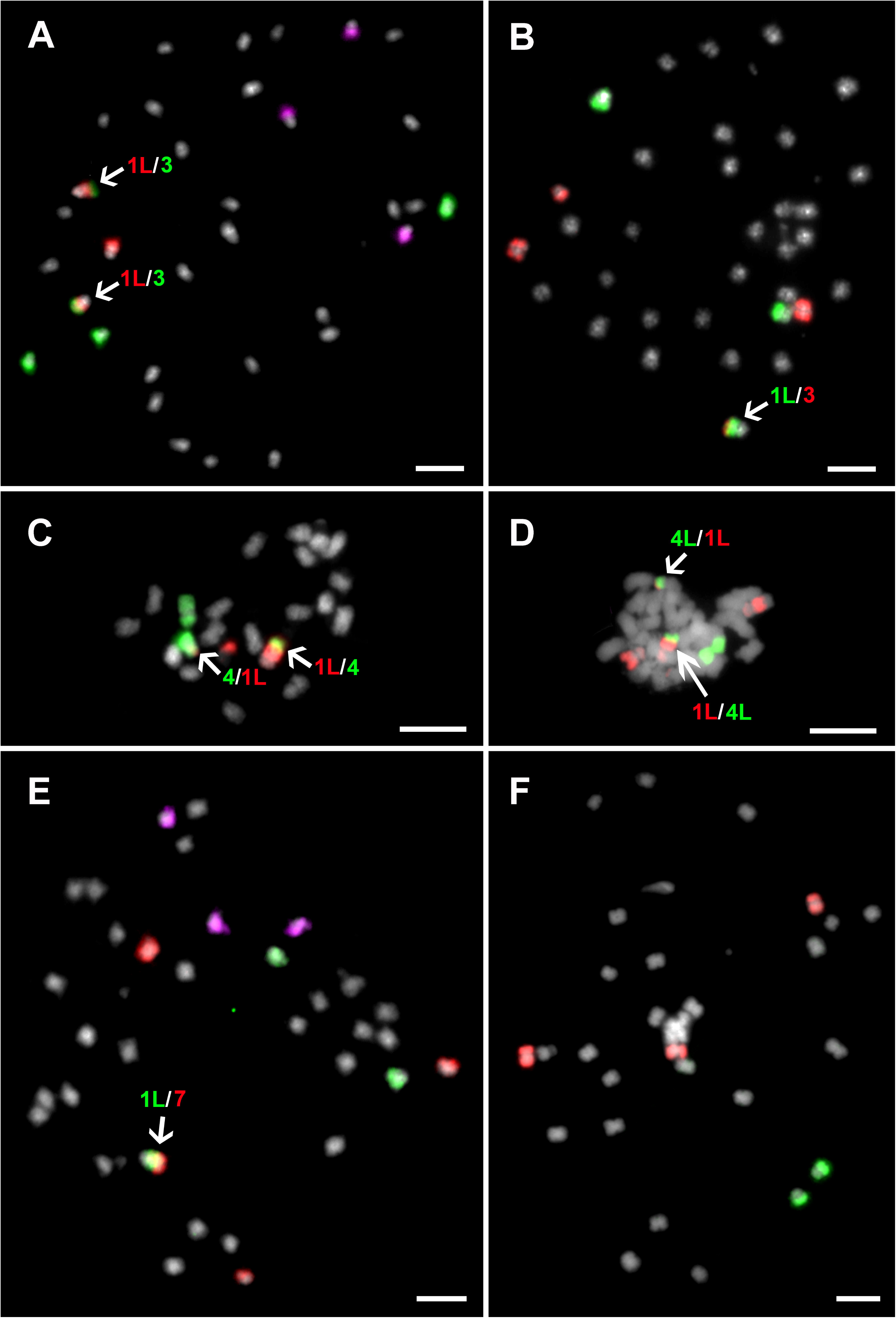
Examples of oligo painting FISH on mitotic metaphase plates of six structurally heterozygous *Musa* accessions: (A) ‘Saba sa Hapon’ (2n=3x=33, ABB genome), probes for long arm of chromosome 1, long arm of chromosome 2 and chromosome 3 were labeled in red, purple and green, respectively; (B) ‘3 Hands Planty’ (2n=3x=33, AAB genome), probes for long arm of chromosome 1 and chromosome 3 were labeled in green and red, respectively; (C) ‘Pisang Lilin’ (2n=2x=22, AA genome), probes for long arm of chromosome 1 and chromosome 4 were labeled in red and green, respectively; (D) ‘Gros Michel’ (2n=3x=33, AAA genome), probes for long arm of chromosome 1 and long arm of chromosome 4 were labeled in red and green, respectively; (E) ‘Gros Michel’ (2n=3x=33, AAA genome), probes for long arm of chromosome 1, long arm of chromosome 2 and chromosome 7 were labeled in green, purple and red, respectively; (F) ‘Amou’ (2n=3x – 1=32, AAB): probes for long arm of chromosome 2 and chromosome 4 were labeled in green and red, respectively. Chromosomes were counterstained with DAPI (light grey pseudocolor). Arrows point translocation chromosomes. Bars = 5 µm.

Two representatives of triploid dessert banana cultivars ‘Cavendish’ and ‘Gros Michel’ with AAA genome constitution displayed identical chromosome structure as assessed by chromosome painting. One of the three chromosome sets was characterized by reciprocal translocation between short segments of the long arms of chromosomes 4 and 1 (Fig. 4D; Suppl. Fig. S1F). It is worth mentioning that the same translocation was identified in Mchare cultivars and in ‘Pisang Lilin’ (Fig. 4C; Suppl. Figs. S1D, S1E). Another chromosome set of ‘Cavendish’ (Fig. 2F) and ‘Gros Michel’ (Fig. 2E) contained a reciprocal translocation between the short arm of chromosome 3 and the long arm of chromosome 8, which was also identified in *M. acuminata* ssp. *zebrina* (Figs. 1B, 2A). In addition to the two translocations, a translocation of the short arm of chromosome 7 to the long arm of chromosome 1 was observed in both accessions of desert banana. The translocation resulted in formation of a small telocentric chromosome consisting of only the long arm of chromosome 7 (Fig. 4E; Suppl. Fig. S1F).

East African Highland bananas (EAHB) represent an important group of triploid cultivars with genome constitution AAA. We have analyzed two accessions of these cooking bananas. Both cultivars (‘Imbogo’ and ‘Kagera’) contained a reciprocal translocation between the short arm of chromosome 3 and the long arm of chromosome 8, which was also observed in *M. acuminata* ssp. *zebrina*. This translocation was identified only in two out of the three chromosome sets (Fig. 2C; Suppl. Figs. S1G, S1H). Karyotype analysis revealed that cultivar ‘Imbogo’ lacked one chromosome and was aneuploid (2n = 32). Chromosome painting facilitated identification of the missing chromosome and suggested the origin of the aneuploid karyotype (Suppl. Figs. S1G, S2), which involved a Robertsonian translocation between chromosomes 7 and 1, giving rise to a recombined chromosome containing long arms of chromosomes 7 and 1. Our observation suggests that short arms of the two chromosomes were lost (Suppl. Figs. S1G, S2). The loss of a putative chromosome comprising two short arms is a common consequence of the Robertsonian translocation (Robertson, 1916).

### Karyotype structure of inter-specific banana clones

Plantains are an important group of starchy type of bananas with AAB genome constitution and originated as hybrids between *M. acuminata* (A genome) and *M. balbisiana* (B genome). Chromosome painting in three cultivars representing two plantain morphotypes - Horn (cultivar ‘3 Hands Planty’) and French type (cultivars ‘Obino l’Ewai’ and ‘Amou’) confirmed the presence of B-genome specific translocation in one chromosome set in all three accessions, i.e. the translocation of a small segment of the long arm of chromosome 3 to the long arm of chromosome 1 (Fig. 4B; Suppl. Figs. S1I, S1J). No other translocation was found in these cultivars. However, the clone ‘Amou’ was found to be aneuploid (2n = 32) as it missed one copy of chromosome 2 (Fig. 4F; Suppl. Fig. S1J).

In agreement with their predicted ABB genome constitution, B genome-specific chromosome translocation was also observed in two chromosome sets in triploid cultivars ‘Pelipita’ and ‘Saba sa Hapon’ (Fig. 4A, Suppl. Fig. S1K). No other translocation was found in these cultivars.

## Discussion

Until recently, cytogenetic analysis in plants was hindered by the lack of available DNA probes suitable for fluorescent painting of individual chromosomes (Schubert *et al*., 2001, Jiang, 2019). The only option was to use pools of BAC clones which were found useful in the plants species with small genomes (e.g. Lysák *et al*., 2001, Idziak *et al*., 2014). The development of oligo painting FISH (Han *et al*., 2015) changed this situation dramatically and it is now possible to label individual plant chromosomes and chromosomal regions in many species (Qu *et al*., 2017, Braz *et al*., 2018, He *et al*., 2018, Machado *et al*., 2018, Albert *et al*., 2019, Bi *et al*., 2020). Chromosome arm-specific oligo painting probes were recently developed also for banana (*Musa* spp.) by Šimoníková *et al*. (2019) who demonstrated the utility of this approach for anchoring DNA pseudomolecules of a reference genome sequence to individual chromosomes *in situ*, and for identification of chromosome translocations.

In this work we used oligo painting FISH for comparative karyotype analysis in a set of *Musa* accessions comprising wild species used in banana breeding programs and economically significant edible cultivars (Suppl. Tab. S1). These experiments revealed chromosomal translocations in subspecies of *M. acuminata* (A genome), their intra-specific hybrids as well as in *M. balbisiana* (B genome) and in inter-specific hybrid clones originating from cross hybridization between *M. acuminata* and *M. balbisiana* (Figure 5). A difference in chromosome structure among *M. acuminata* subspecies was suggested earlier by Shepherd *et al*. (1999) who identified seven translocation groups in *M. acuminata* based on chromosome pairing during meiosis. An independent confirmation of this classification was the observation of segregation distortion during genetic mapping in inter-subspecific hybrids of *M. acuminata* (Fauré *et al*., 1993, Hippolyte *et al*., 2010, Mbanjo *et al*., 2012, Noumbissie *et al*., 2016).

**Figure 5.**
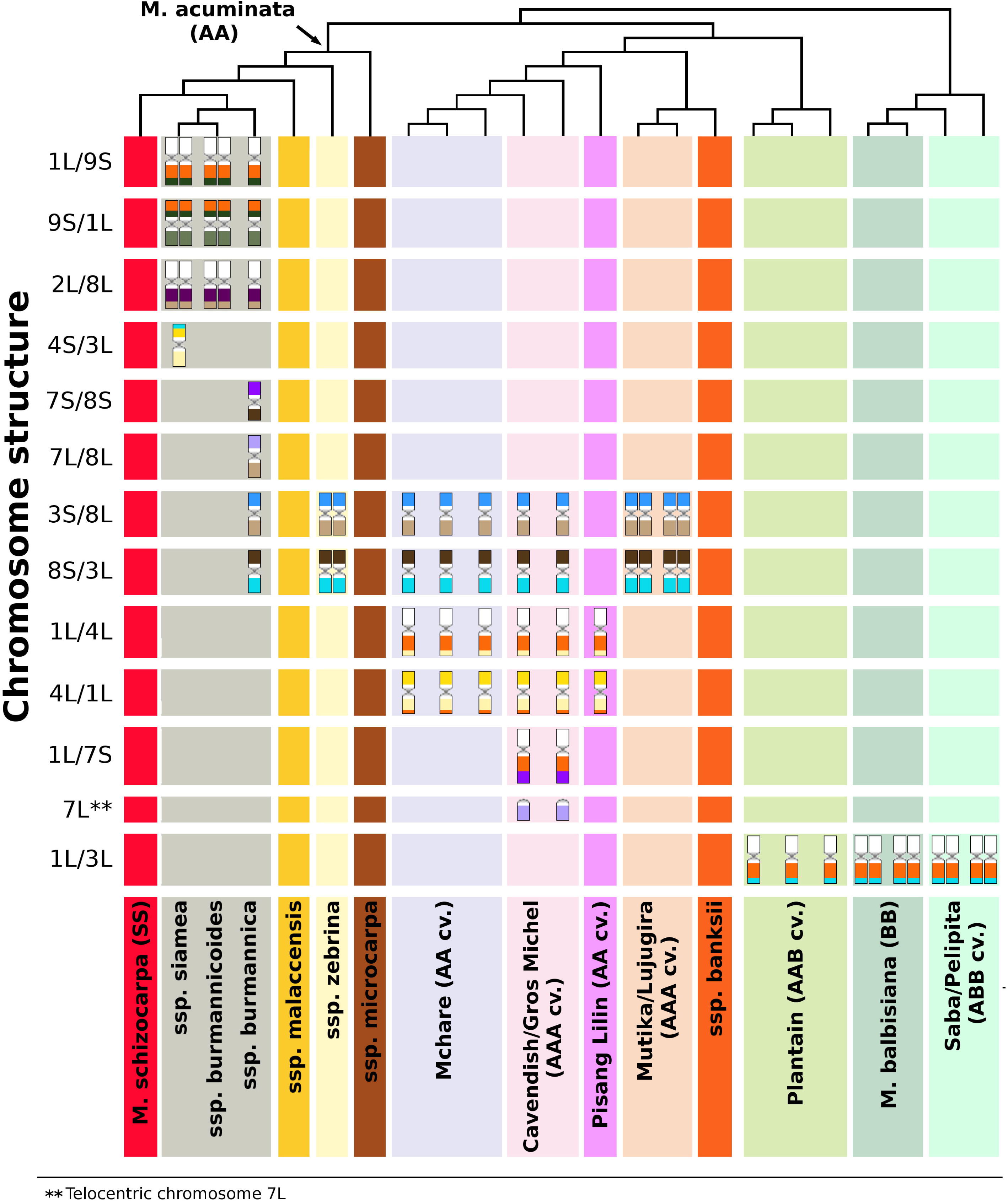
Overview of common translocation events revealed by oligo painting FISH in *Musa*: Diversity tree, constructed using SSR markers according to Christelová *et al*. (2017), shows the relationships among *Musa* species, subspecies and hybrid clones. Lineages of closely related accessions and groups of edible banana clones are highlighted in different colors. Individual chromosome structures (displayed as chromosome schemes) are depicted in rows, and their number correspond to the number of chromosomes bearing the rearrangement in the nuclear genome in somatic cell lines (2n).

### Structural genome variation in diploid *M. acuminata*

In this work we analyzed one representative of each of six subspecies of *M. acuminata*. We cannot exclude differences in chromosome structure within individual subspecies. However, given the large genetic homogeneity of the six subspecies as clearly demonstrated by molecular markers (Carreel *et al*., 2002, Perrier *et al*., 2009, Hřibová *et al*., 2011, Christelová *et al*., 2017, Němečková *et al*., 2018, Dupouy *et al*., 2019), this does not seem probable. We observed a conserved genome structure in *M. acuminata* ssp. *banksii* and *M. acuminata* ssp. *microcarpa* which did not contain any translocation chromosome when compared to the reference genome of *M. acuminata* ssp. *malaccensis* ‘DH Pahang’ (Martin *et al*., 2016). The genome structure shared by the three subspecies was also observed in *M. schizocarpa* (Šimoníková *et al*., 2019) and corresponds to the standard translocation (ST) group as defined by Shepherd (1999).

Other chromosome translocation group defined by Shepherd (1999), the Northern Malayan group (NM), is characteristic for *M. acuminata* ssp. *malaccensis* and some AA cultivars, including ‘Pisang Lilin’. Our results revealed a reciprocal translocation between chromosomes 1 and 4 in one chromosome set of ‘Pisang Lilin’, thus confirming Shepherd’s characterization of this clone as heterozygous having ST x NM genome structure. Before the *Musa* genome sequence became available, Hippolyte *et al*. (2010) assumed a presence of a duplication of distal region of chromosome 1 on chromosome 4 in this clone based on comparison of high dense genetic maps. Taking the advantage of the availability of reference genome sequence and after resequencing genomes of a set of *Musa* species, Martin *et al*. (2017) described heterozygous reciprocal translocation between chromosomes 1 and 4, involving 3 Mb of long arm of chromosome arm 1 and 10 Mb segment of long arm of chromosome arm 4 in *M. acuminata* ssp. *malaccensis*. Further experiments indicated preferential transmission of the translocation to the progeny, and its frequent presence in triploid banana cultivars (Martin *et al*., 2017). Yet, it is not clear if the reciprocal translocation between chromosomes 1 and 4, which we observed in the heterozygous state only, originated in ssp. *malaccensis*, or if it was transmitted to genomes of some *malaccensis* accessions by ancient hybridization events (Martin *et al*., 2017). Three phylogenetically closely related subspecies *M. acuminata* ssp. *burmannica, M. acuminata* ssp. *burmannicoides* and *M. acuminata* ssp. *siamea*, which have similar phenotype and geographic distribution (Simmonds, 1962, Perrier *et al*., 2009) share a translocation of a part of the long arm of chromosome 8 to the long arm of chromosome 2, and a reciprocal translocation between chromosomes 1 and 9. These translocations were identified recently by Dupouy *et al*. (2019) after mapping mate-paired Illumina sequence reads to the reference genome of ‘DH Pahang’. The authors estimated the size of translocated region of chromosome 8 to chromosome 2 to be 7.2 Mb, while the size of the distal region of chromosome 2, which was found translocated to chromosome 8 in wild diploid clone ‘Calcutta 4’ (ssp. *burmannicoides*) was estimated to be 240 kb (Dupouy *et al*., 2019). The size of the translocated regions of chromosomes 1 and 9 was estimated to be 20.8 Mb and 11.6 Mb respectively (Dupouy *et al*., 2019). Using oligo painting, we did not detect the 240 kb distal region of chromosome 2 translocated to chromosome 8, and this may reflect the limitation in the sensitivity of whole chromosome arm oligo painting.

The shared translocations in all three subspecies of *M. acuminata* (*burmannicoides, burmannica* and *siamea*) support their close phylogenetic relationship as proposed by Shepherd (1999) and later verified by molecular studies (Carreel *et al*., 2002, Perrier *et al*., 2009, Hřibová *et al*., 2011, Christelová *et al*., 2017, Němečková *et al*., 2018). Dupouy *et al*. (2019) coined the idea of a genetically uniform *burmannica* group. However, our data indicate a more complicated evolution of the three genotypes recognized as representatives of different *acuminata* subspecies. First, the characteristic translocations between chromosomes 2 and 8, and 1 and 9 were detected only on one chromosome set in *burmannica* ‘Tavoy’, as compared to *M. acuminata* ‘Calcutta 4’ (ssp. *burmannicoides*) and ‘Pa Rayong’ (ssp. *siamea*). Second, we observed two subspecies-specific translocations in ssp. *burmannica* only in one chromosome set, and we detected additional subspecies-specific translocation in *M. acuminata* ssp. *siamea* only in one chromosome set, indicating its hybrid origin. Based on these results we hypothesize that ssp. *burmannicoides* could be a progenitor of the clones characterized by structural chromosome heterozygosity. Divergence in genome structure between the three subspecies (*burmannica, burmannicoides* and *siamea*) was demonstrated also by Shepherd (1999), who classified some *burmannica* and *siamea* accessions as Northern 2 translocation group of *Musa*, differing from the Northern 1 group (*burmannicoides* and other *siamea* accessions) by one additional translocation.

We observed subspecies-specific translocations also in *M. acuminata* ssp. *zebrina* ‘Maia Oa’. In this case, chromosome painting revealed a Robertsonian translocation between chromosomes 3 and 8. Interestingly, Dupouy *et al*. (2019) failed to detect this translocation after sequencing mate-pair libraries using Illumina technology. The discrepancy may point to the limitation of the sequencing approach to identify translocations arising by a breakage of (peri)centromeric regions. As these regions comprise mainly DNA repeats, they may not be assembled properly in a reference genome, thus preventing their identification by sequencing. In fact, this problem may also be encountered if subspecies-specific genome regions are absent in the reference genome sequence. We revealed the *zebrina*-type translocation (a reciprocal translocation between chromosomes 3 and 8) also in all three analyzed cultivars of diploid Mchare banana. The presence of a translocation between long arm of chromosome 1 and long arm of chromosome 4 on one chromosome set of Mchare indicates a hybrid origin of Mchare, with ssp. *zebrina* being one of the progenitors of this banana group. This agrees with the results obtained by genotyping using molecular markers (Christelová *et al*., 2017). Most recently, complex hybridization scheme of Mchare bananas was supported also by application of transcriptomic data for identification of specific SNPs in 23 *Musa* species and edible cultivars (Martin *et al*., 2020).

### Genome structure and origin of cultivated triploid *Musa* clones

Plantains are an important group of triploid starchy bananas with AAB genome constitution, which originated after hybridization between *M. acuminata* and *M. balbisiana*. As expected, chromosome painting in ‘3 Hands Planty’, ‘Amou’ and ‘Obino l’Ewai’ cultivars revealed B-genome specific translocation of 8 Mb segment from the long arm of chromosome 3 to the long arm of chromosome 1. Unlike the B-genome chromosome set, the two A-genome chromosome sets of plantains lacked any detectable translocation. Genotyping using molecular markers revealed that the A genomes of the plantain group are related to *M. acuminata* ssp. *banksii* (Horry, 1989, Lebot *et al*., 1993, Carreel *et al*., 2002, Kagy *et al*., 2016). This is in line with the absence of chromosome translocations we observed in *M. acuminata* ssp. *banksii*.

Triploid cultivar ‘Pisang Awak’, a representative of the ABB group, is believed to contain one A genome chromosome set closely related to *M. acuminata* ssp. *malaccensis* (Perrier *et al*., 2009). However, after SSR genotyping Christelová *et al*. (2017) found, that some ABB clones from Saba and Bluggoe-Monthan groups clustered together with the representatives of Pacific banana Maia Maoli / Popoulu (AAB). These results point to *M. acuminata* ssp. *banksii* as their most probable progenitor. Unfortunately, in this case chromosome painting did not bring useful hints on the nature of the A subgenomes in these inter-specific hybrids, as *M. acuminata* ssp. *banksii* and ssp. *malaccensis* representatives do not differ in the presence of specific chromosome translocation.

The presence of B genome-specific translocation of a small region of long arm of chromosome 3 to the long arm of chromosome 1 (Šimoníková *et al*., 2019), observed in all *M. balbisiana* accessions and their inter-specific hybrids with *M. acuminata*, seems to be a useful cytogenetic landmark of the presence of B sub-genome. In our study, the number of chromosome sets containing B genome specific translocation agreed with the predicted genomic constitution (AAB or ABB) of the hybrids. Clearly, one B genome-specific landmark is not sufficient for analysis of complete genome structure of inter-specific hybrids at the cytogenetic level. Further work is needed to identify additional cytogenetic landmarks to uncover the complexities of genome evolution after inter-specific hybridization in *Musa*. Recently, Baurens *et al*. (2019) analyzed genome composition of banana inter-specific hybrid clones using whole genome sequencing strategies followed by bioinformatic analysis based on A- and B-genome specific SNPs calling and they also detected the B genome-specific translocation in inter-specific hybrids.

Chromosome painting confirmed a small genetic difference between triploid clones ‘Gros Michel’ and ‘Cavendish’ (AAA genomes) as previously determined by various molecular studies (Raboin *et al*., 2005, Christelová *et al*., 2017). Both clones share the same reciprocal translocation between long arms of chromosomes 1 and 4 in one chromosome set. Interestingly, also diploid Mchare cultivars contain the same translocation in one chromosome set. The presence of the translocation was identified also by sequencing genomic DNA both in ‘Cavendish’ and ‘Gros Michel’, as well as in Mchare banana ‘Akondro Mainty’ (Martin *et al*., 2017). These observations confirm close genetic relationship between both groups of edible bananas as noted previously (Raboin *et al*., 2005, Perrier *et al*., 2009). According to Martin *et al*. (2017, 2020), 2n gamete donor, which contributed to the origin of dessert banana clones with AAA genomes, including ‘Cavendish’ and ‘Gros Michel’, belongs to the Mchare (Mlali) sub-group. The genome of this ancient sub-group, which probably originated somewhere around Java, Borneo and New Guinea, but today is only found in East Africa, is based on *zebrina / microcarpa* and *banksii* subspecies (Perrier *et al*., 2009).

The third chromosome set in triploid ‘Cavendish’ and ‘Gros Michel’ contains a reciprocal translocation between chromosomes 3 and 8, which was detected by oligo painting FISH in the diploid *M. acuminata* ssp. *zebrina*. Our observations indicate that heterozygous representatives of *M. acuminata* ssp. *malaccensis* and *M. acuminata* spp. *zebrina* contributed to the origin of ‘Cavendish’ and ‘Gros Michel’ as their ancestors as suggested earlier (Perrier *et al*., 2009, Hippolyte *et al*., 2012, Christelová *et al*., 2017). According to Perrier *et al*. (2009), *M. acuminata* ssp. *banksii* was one of the two progenitors of ‘Cavendish’ / ‘Gros Michel’ group of cultivars. However, using chromosome arm-specific oligo painting, we observed a translocation of the short arm of chromosome 7 to the long arm of chromosome 1, resulting in a small telocentric chromosome made only of the long arm of chromosome 7, which was, up to now, identified only in these cultivars. The translocation, which gave arise to the small telocentric chromosome, could be a result of processes accompanying the evolution of this group of triploid AAA cultivars. Alternatively, another wild diploid clone, possibly structurally heterozygous, was involved in the origin of ‘Cavendish’ / ‘Gros Michel’ bananas.

We observed reciprocal translocation between chromosomes 3 and 8, which is typical for *M. acuminata* ssp. *zebrina*, in the economically important group of triploid East African Highland bananas (EAHB) with AAA genome constitution. An important role of *M. acuminata* ssp. *zebrina* and *M. acuminata* ssp. *banksii*, as the most probable progenitors of EAHB, was suggested previously (Carreel *et al*., 2002, Li *et al*., 2013, Kitavi *et al*., 2016, Christelová *et al*., 2017, Němečková *et al*., 2018, Martin *et al*., 2020). Our results, which indicate that EAHB contained two chromosome sets from ssp. *zebrina* and one chromosome set from ssp. *banksii*, point to the most probable origin of EAHB. Hybridization between *M. acuminata* ssp. *zebrina* and ssp. *banksii* could give arise to intra-specific diploid hybrid with a reduced fertility. Triploid EAHB cultivars then could originate by backcross of the intra-specific hybrid (a donor of non-reduced gamete) with *M. acuminata* ssp. *zebrina*, or with another diploid, most probably a hybrid of *M. acuminata* ssp. *zebrina*. Moreover, phylogenetic analysis of Němečková *et al*. (2018) surprisingly revealed also possible contribution of *M. schizocarpa* to EAHB formation, thus indicating a more complicated origin. Further investigation is needed, and the availability of EAHB genome sequence in particular, to shed more light on the origin and evolution of these triploid clones.

### The origin of aneuploidy

To date, aneuploidy in *Musa* has been identified by chromosome counting (Sandoval *et al*., 1996, Shepherd and Da Silva, 1996, Bartoš *et al*., 2005, Čížková *et al*., 2013, Čížková *et al*., 2015, Němečková *et al*., 2018). Although this approach is laborious and low throughput, it cannot be replaced by flow cytometric estimation of nuclear DNA amounts because of the differences in genome size between *Musa* species, sub-species and their hybrids (e.g. Čížková *et al*., 2013, Christelová *et al*., 2017, Němečková *et al*., 2018). A more laborious approach to achieve high resolution flow cytometry as used by Roux *et al*. (2003) is too slow and laborious to be practical. Thus, traditional chromosome counting (Sandoval *et al*., 1996, Shepherd and Da Silva, 1996, Bartoš *et al*., 2005, Čížková *et al*., 2013, Čížková *et al*., 2015, Němečková *et al*., 2018) remains the most reliable approach. Obviously, it is not suitable to identify the chromosome(s) involved in aneuploidy and the origin of the aberrations.

Here we employed chromosome painting to shed light on the nature of aneuploids among the triploid *Musa* accessions. One of the aneuploid clones was identified in plantain ‘Amou’, in which one copy of chromosome 2 was lost. The origin of aneuploidy in clone ‘Imbogo’ (AAA genome), a representative of EAHB, involved structural chromosome changes involving breakage of chromosomes 1 and 7 in centromeric regions, followed by fusion of long arms of chromosomes 1 and 7 and subsequent loss of short arms of both chromosomes (Suppl. Fig. S2). It needs to be noted that these plants were obtained from the International Musa Germplasm Transit Centre (ITC, Leuven, Belgium), where the clones are stored *in vitro*. The loss of whole chromosome in plantain ‘Amou’ could occur during long-term culture.

To conclude, the application of oligo painting FISH improved the knowledge on genomes of cultivated banana and their wild relatives at chromosomal level. For the first time, a comparative molecular cytogenetic analysis of twenty representatives of the Eumusa section of genus *Musa*, including accessions commonly used in banana breeding, was performed using chromosome painting. However, as only a single accession from each of the six subspecies of *M. acuminata* was used in this study, our results will need to be confirmed by analyzing more representatives from each taxon. The identification of chromosome translocations pointed to particular *Musa* subspecies as putative parents of cultivated clones and provided an independent support for hypotheses on their origin. While we have unambiguously identified a range of translocations, a precise determination of breakpoint positions will have to be done using a long read DNA sequencing technology (Sedlazeck *et al*., 2018, Hu *et al*., 2020, Soto *et al*., 2020).

The discrepancies in genome structure of banana diploids observed in our study and published data then point to alternative scenarios on the origin of the important crop. The observation on structural chromosome heterozygosity confirmed the hybrid origin of cultivated banana and some of the wild diploid accessions which were described as individual subspecies, and informs breeders on possible causes of reduced fertility. The knowledge on genome structure at chromosomal level and characterization structural chromosome heterozygosity will aid breeders in selecting parents for improvement programs.

## Abbreviations

BAC: bacterial artificial chromosome
DH Pahang: doubled haploid Pahang
EAHB: East African Highland banana
FISH: fluorescence *in situ* hybridization
FITC: fluorescein isothiocyanate
NM group: Northern Malayan group
SNP: single-nucleotide polymoprhism
*spp*.: species
*ssp*.: subspecies
SSR: simple sequence repeat
ST group: standard translocation group
UPGMA: unweighted pair group method with arithmetic mean

## Acknowledgements

We thank Dr. Ines van den Houwe for providing the plant material. This work was supported by the Czech Science Foundation (award No. 19-20303S) and by the Ministry of Education, Youth and Sports of the Czech Republic (Program INTER-EXCELLENCE, INTER-TRANSFER, grant award LTT19, and ERDF project “Plants as a tool for sustainable global development” No. CZ.02.1.01/0.0/0.0/16_019/0000827). The computing was supported by the National Grid Infrastructure MetaCentrum (grant No. LM2010005 under the program Projects of Large Infrastructure for Research, Development, and Innovations). The authors thank all donors who supported this work also through their contributions to the CGIAR Fund (http://www.cgiar.org/who-we-are/cgiar-fund/fund-donors-2/), and in particular to the CGIAR Research Program Roots, Tubers and Bananas (RTB-CRP).

## Author contributions

EH and JD conceived the project. DŠ, AN and JČ conducted the study and processed the data. AB and RS provided the banana materials. DŠ and EH wrote the manuscript. EH, DŠ, AB, RS and JD discussed the results and contributed to manuscript writing. All authors have read and approved the final manuscript.

## Data statement

Data supporting the findings of this work are available within the paper and its Supporting Information files. All other data generated and analyzed during the current study are available from the corresponding author upon reasonable request.

## Supplementary data

**Supplementary Figure S1:** Idiograms of *Musa* accessions. (A) *M. acuminata* ssp. *siamea* ‘Pa Rayong’ (genome AA); (B) *M. acuminata* ssp. *burmannica* ‘Tavoy’ (genome AA); (C) *M. balbisiana* ‘Pisang Klutuk Wulung’ (genome BB); (D) *M. acuminata* ‘Pisang Lilin’ (genome AA); (E) *M. acuminata* subgr. Mchare, clones ‘Huti white’, ‘Huti (Shumba nyeelu)’ and ‘Ndyali’ (genomes AA); (F) triploid clones ‘Gros Michel’ and ‘Poyo’ (genomes AAA); (G) East African Highland Banana (EAHB) clone ‘Imbogo’ (genome AAA); (H) EAHB clone ‘Kagera’ (genome AAA); (I) plantains ‘3 Hands Planty’ and ‘Obino l’Ewai’ (genomes AAB); (J) aneuploid plantain clone ‘Amou’; (K) triploid clones ‘Pelipita’ and ‘Saba sa Hapon’ (genomes ABB). Short arms of the chromosomes are on the top and the long arms on the bottom in all idiograms. For better orientation, translocated parts of the chromosomes contain extra labels, which include chromosome number and chromosome arm which was involved in the rearrangement.

**Supplementary Figure S2:** Robertsonian translocation as a mechanism leading to structural chromosome change (formation of a translocation chromosome 7L.1L) and aneuploidy (loss of the short arms of chromosomes 1 and 7) in East African Highland Banana clone ‘Imbogo’.

**Supplementary Figure S3**: Anchoring of chromosomes to *M. acuminata* ‘DH Pahang’ pseudomolecules which were used to develop oligo painting probes. Note that in the reference genome assembly, pseudomolecules 1, 6, 7 are oriented inversely to the traditional way karyotypes are presented, where the short arms are on the top.

**Supplementary Table S1:** Comparison of genome structure in selected *Musa* accessions compared to the *Musa* reference genome sequence (*M. acuminata* ssp. *malaccensis* ‘DH Pahang’).

