## Supplementary Figure S1 for "Chromosome painting in cultivated banana and their wild relatives (*Musa* spp.) reveals differences in chromosome structure"

A

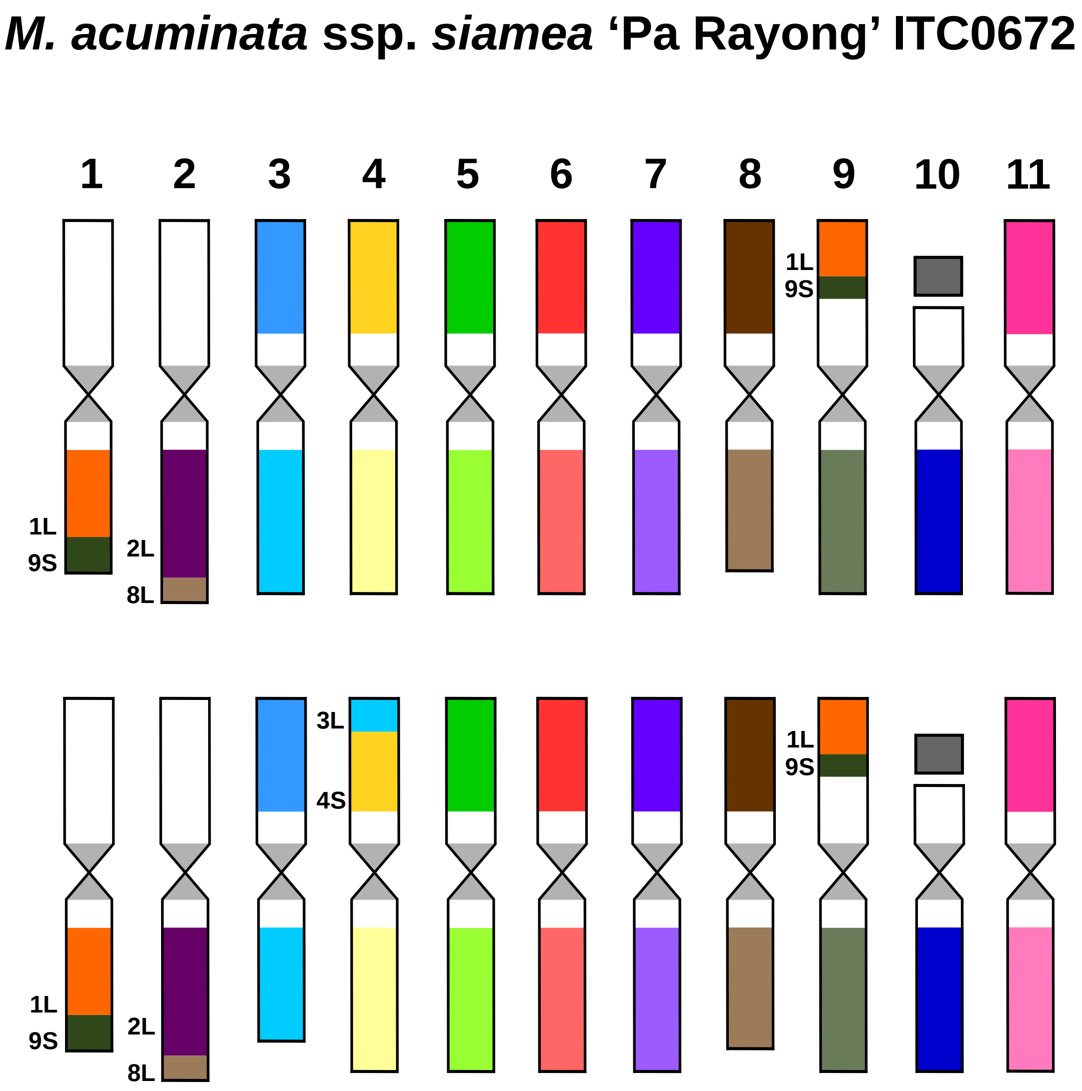

M. acuminata ssp. siamea 'Pa Rayong' ITC0672
1
2
3
4
5
6
7
8
9
10
11
1L
9S
1L
9S
2L
8L
3L
4S
1L
9S
1L
9S
2L
8L

B

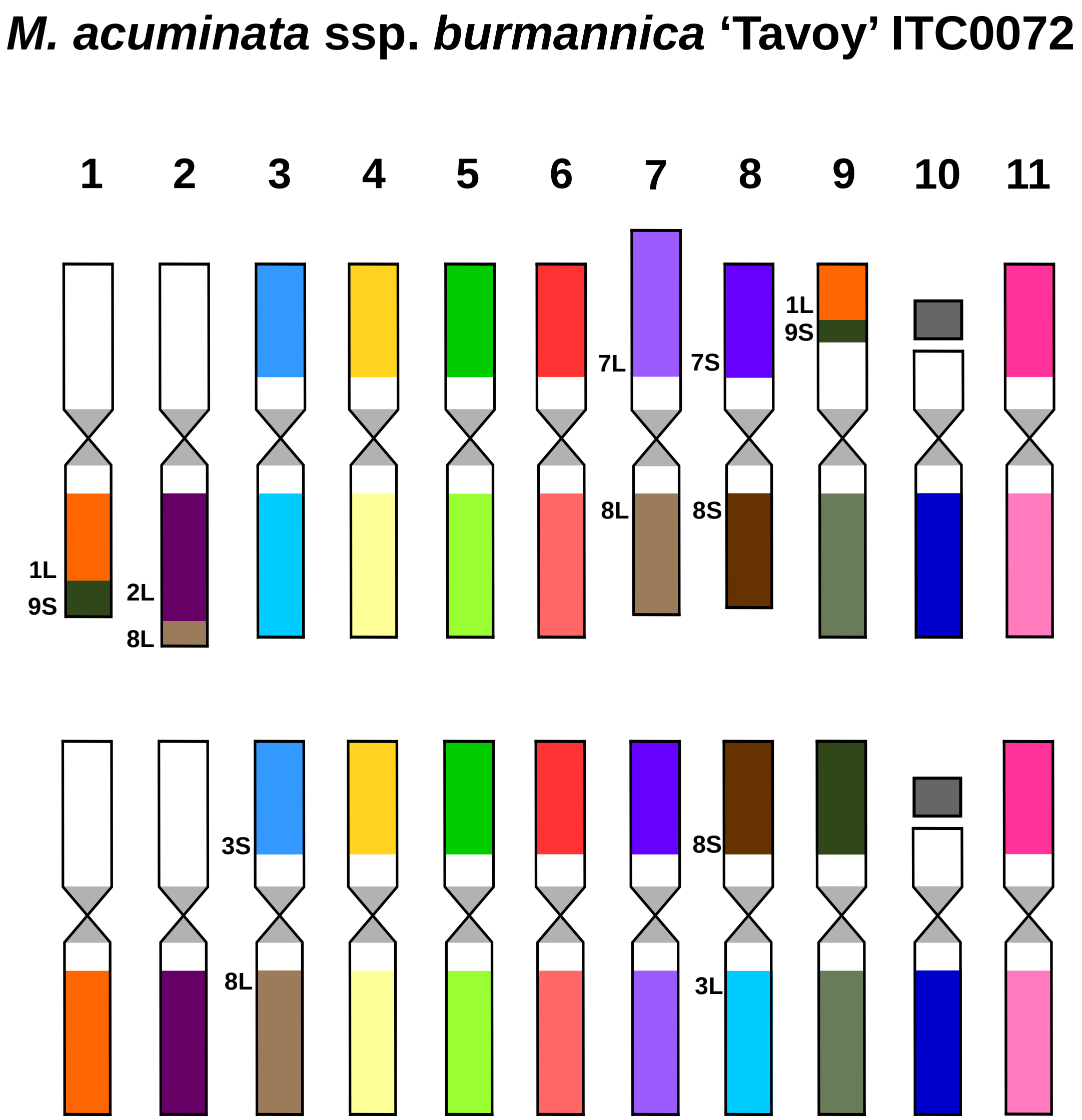

*M. acuminata* ssp. *burmannica* 'Tavoy' ITC0072
1
2
3
4
5
6
7
8
9
10
11
1L
9S
2L
8L
7L
8L
7S
8S
1L
9S
3S
8L
8S
3L

C

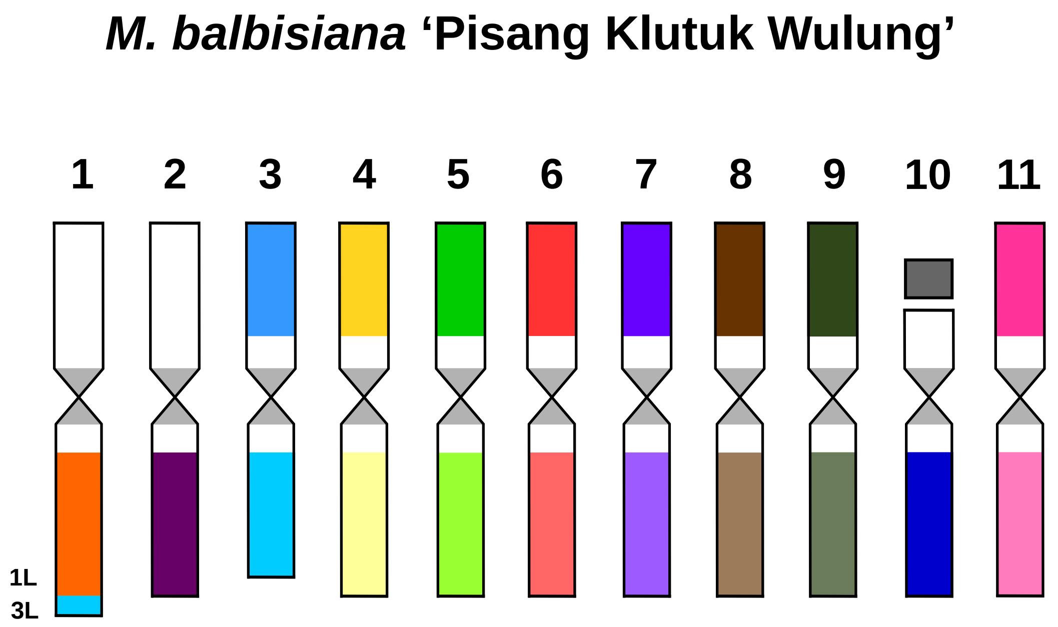

D

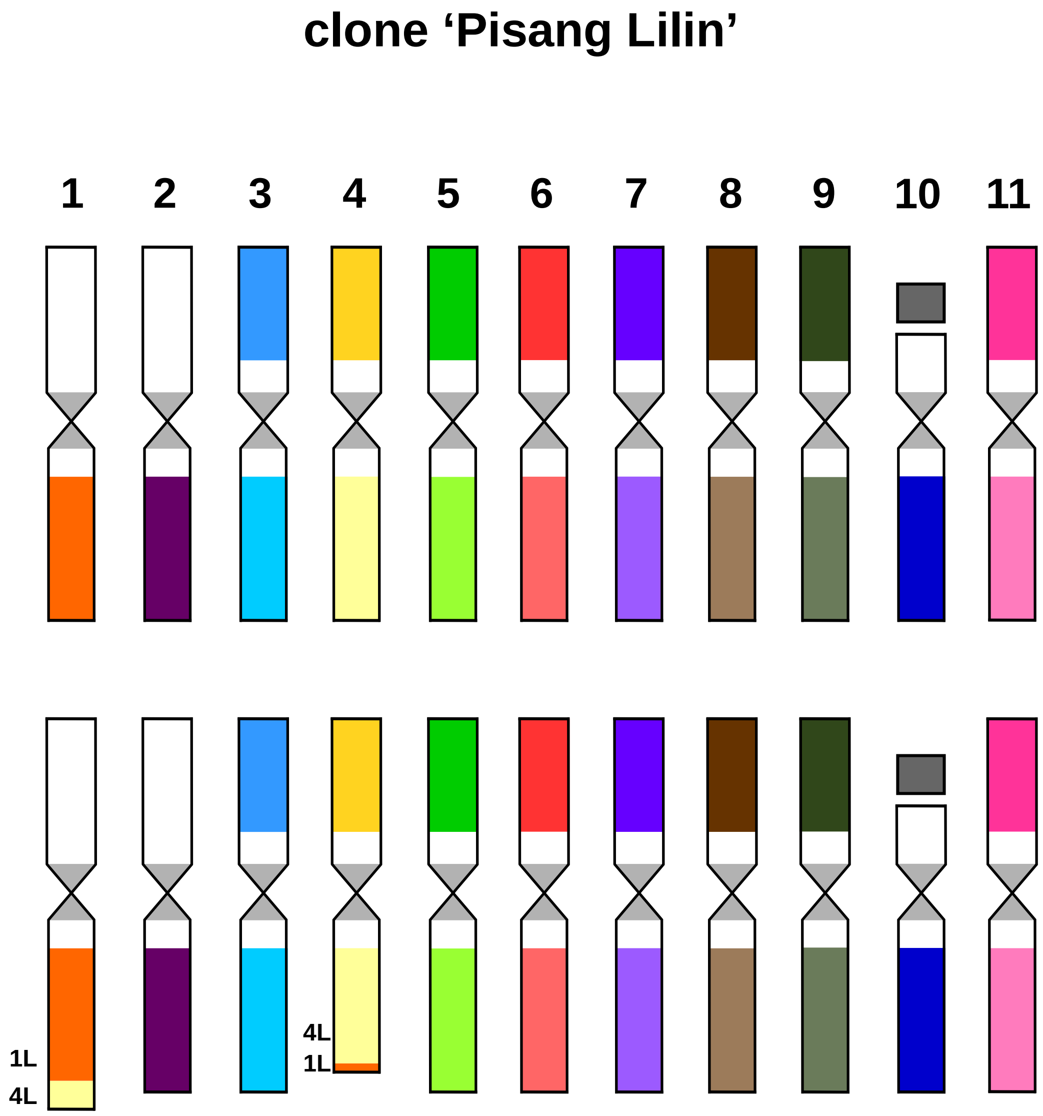

E

### Mchare clones 'Huti white', 'Huti (Shumba nyeelu)' ITC1452, 'Ndyali' ITC1552

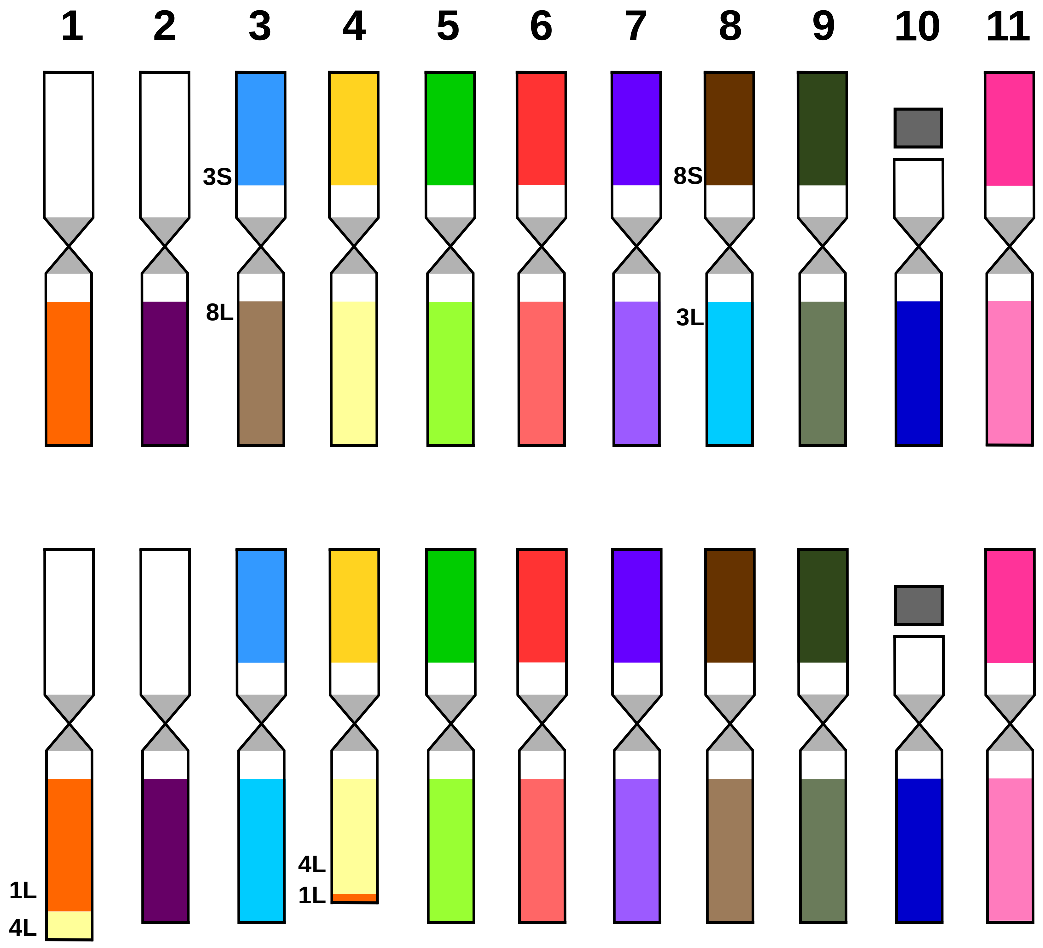

F

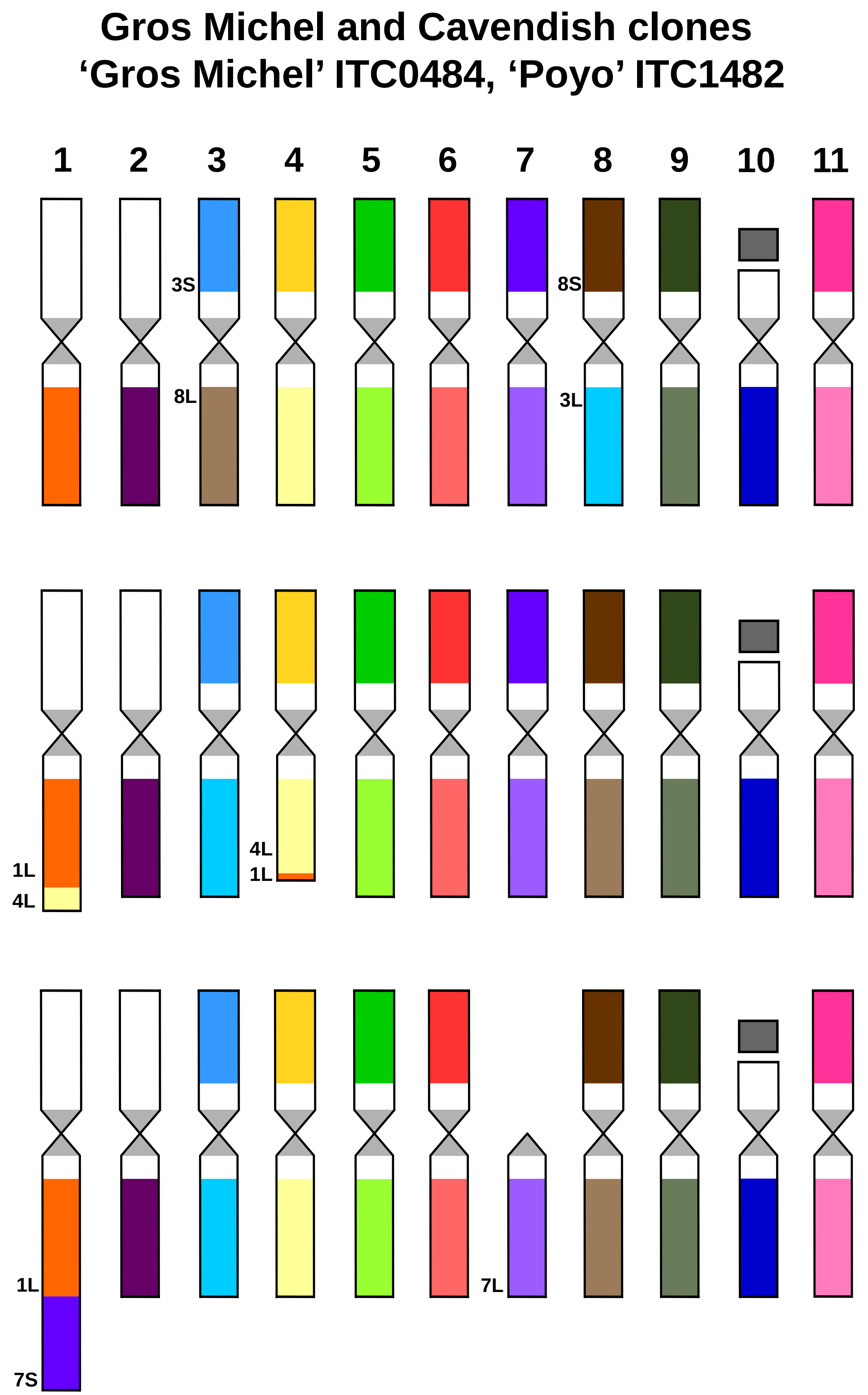

Gros Michel and Cavendish clones
‘Gros Michel’ ITC0484, ‘Poyo’ ITC1482
1
2
3
4
5
6
7
8
9
10
11
3S
8L
8S
3L
1L
4L
4L
1L
1L
7S
7L

G

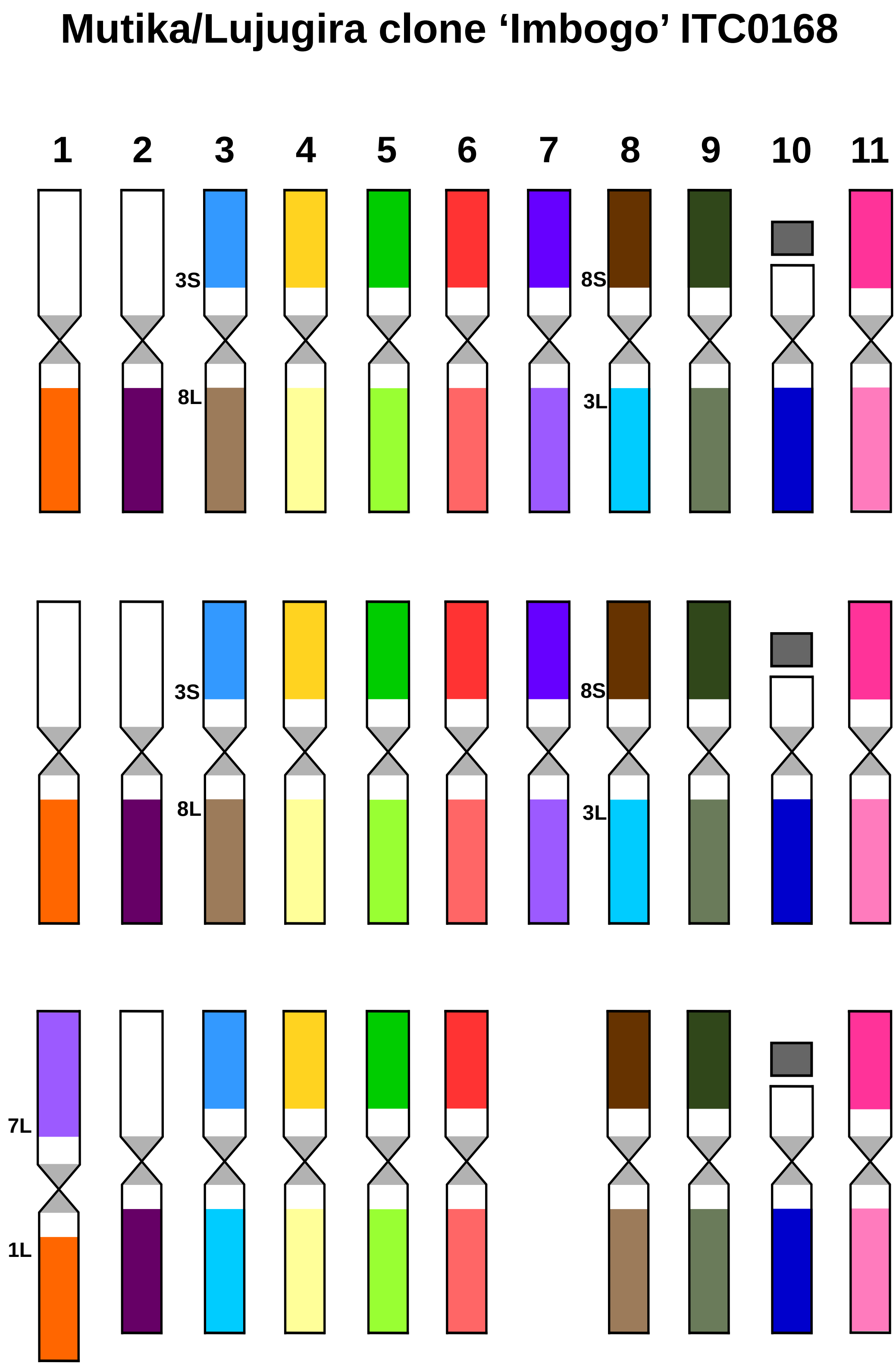

Mutika/Lujugira clone 'Imbogo' ITC0168
1
2
3
4
5
6
7
8
9
10
11
3S
8L
8S
3L
3S
8L
8S
3L
7L
1L

H

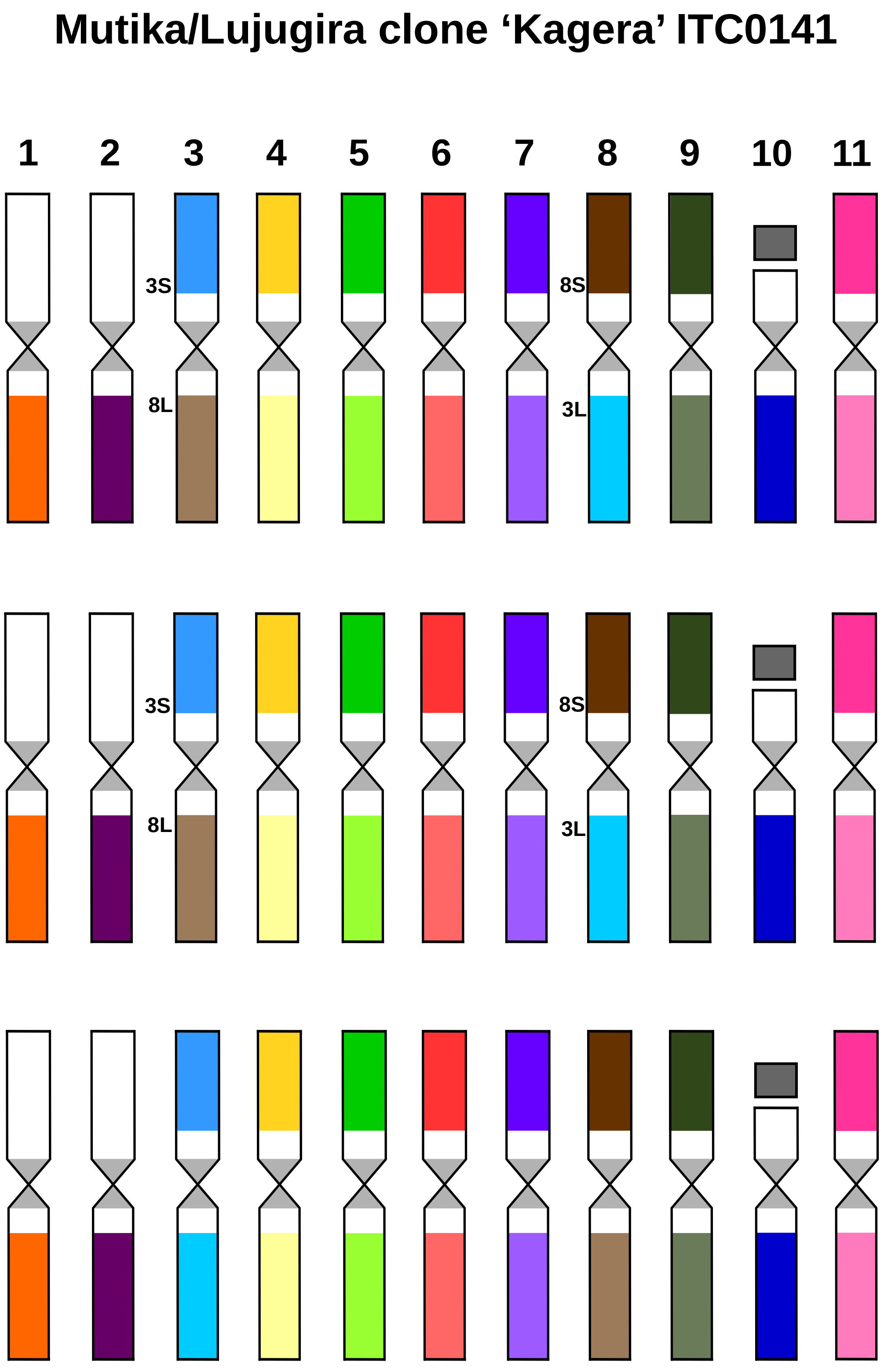
Mutika/Lujugira clone 'Kagera' ITC0141
1
2
3
4
5
6
7
8
9
10
11
3S
8L
8S
3L
3S
8L
8S
3L

I

### Plantain clones '3 Hands Planty' ITC1132, 'Obino l'Ewai' ITC0109

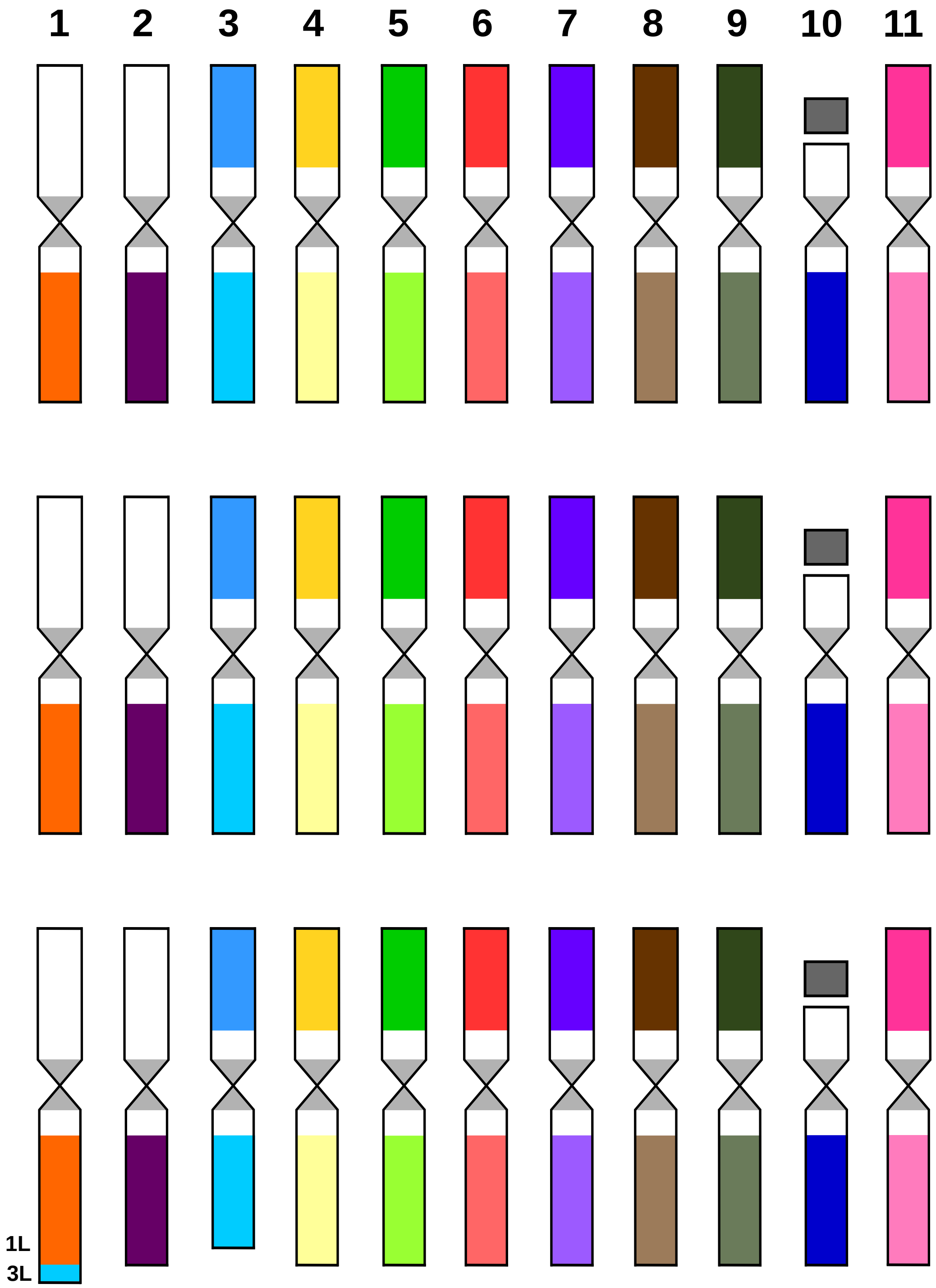

J

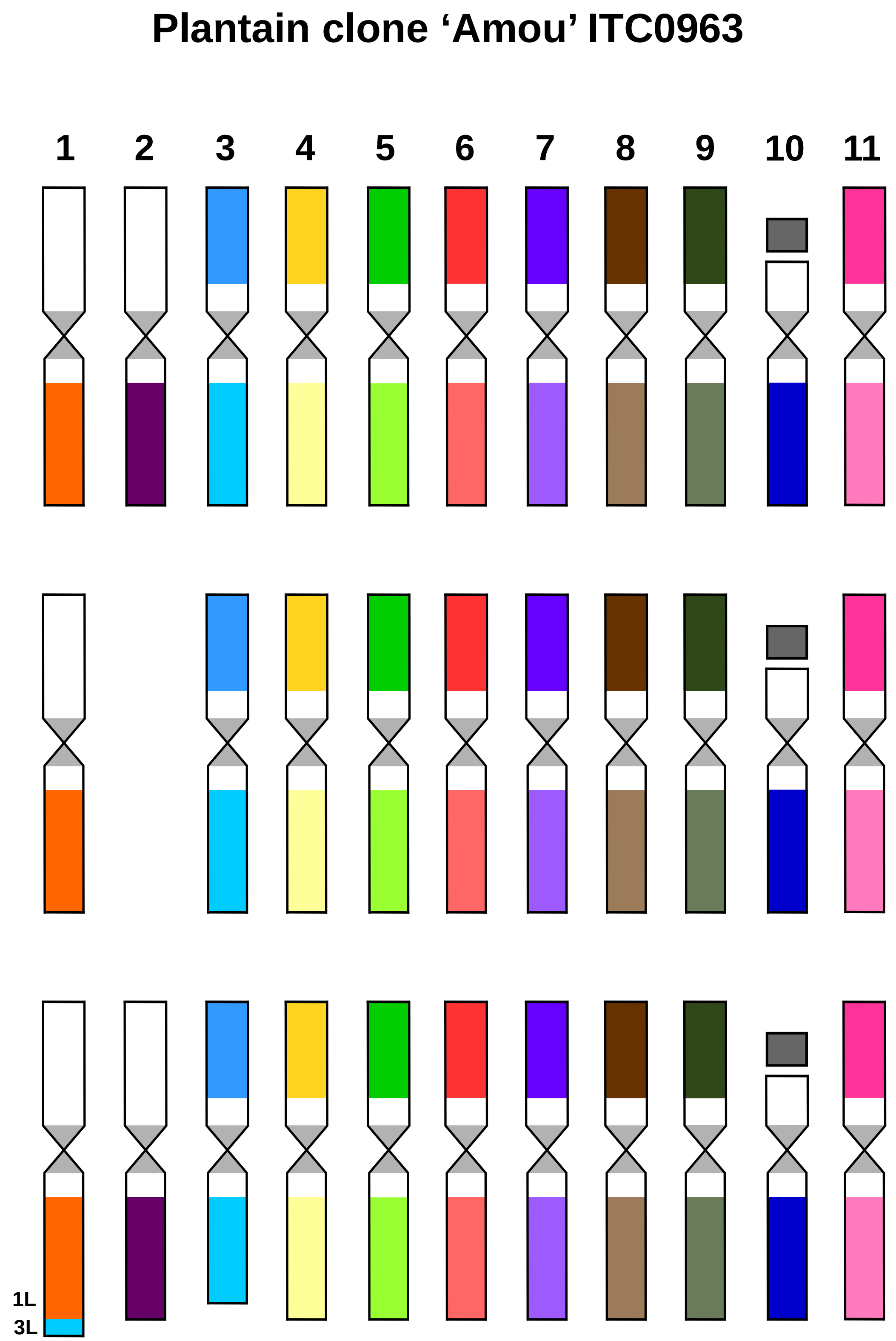

K

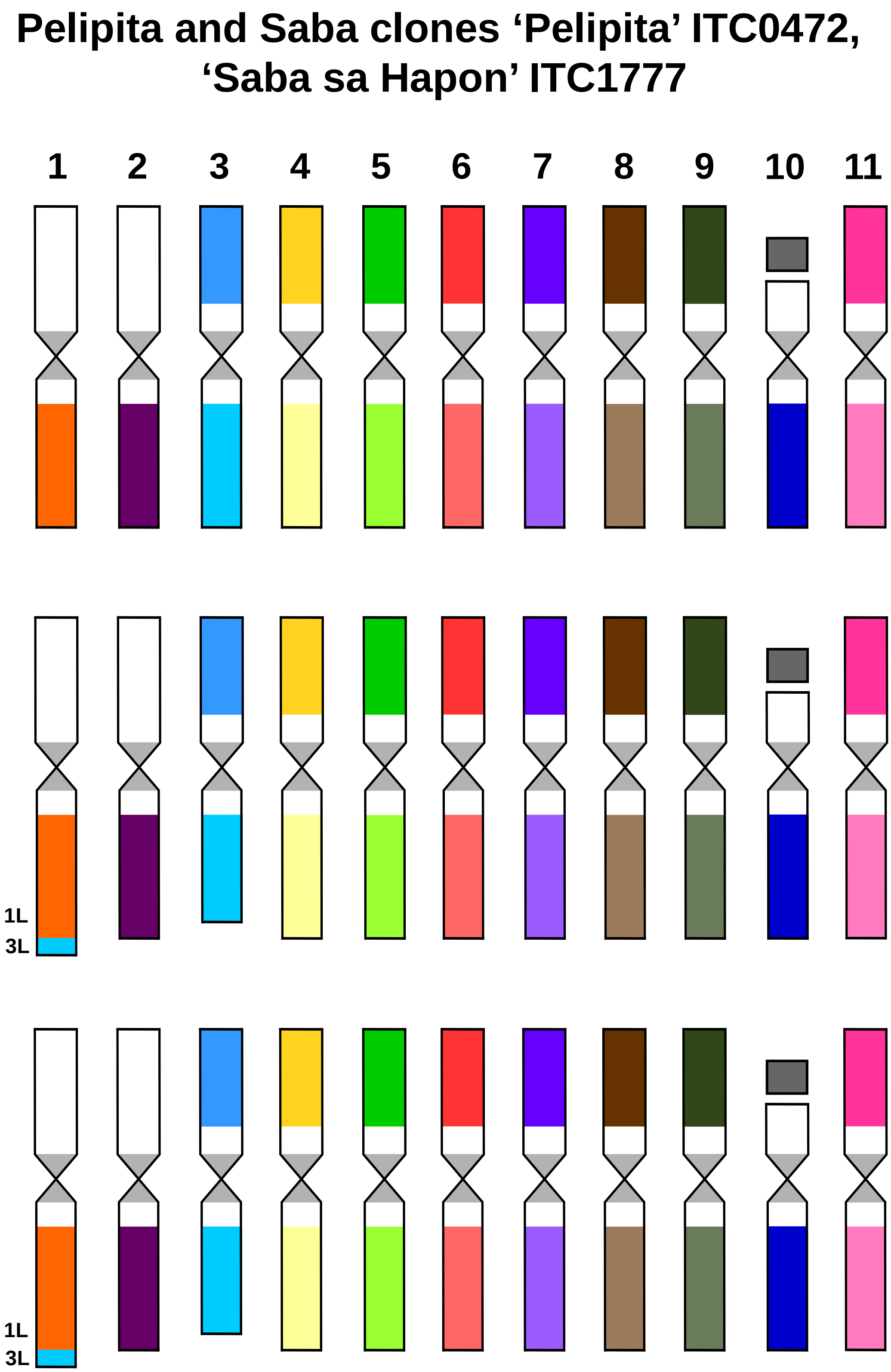
Pelipita and Saba clones 'Pelipita' ITC0472,
'Saba sa Hapon' ITC1777
1
2
3
4
5
6
7
8
9
10
11
1L
3L
1L
3L
