## Supplementary figures and images for "Chromosome painting in cultivated banana and their wild relatives (*Musa* spp.) reveals differences in chromosome structure"

### Supplementary Figure S2

**Supplementary Figure S2.**

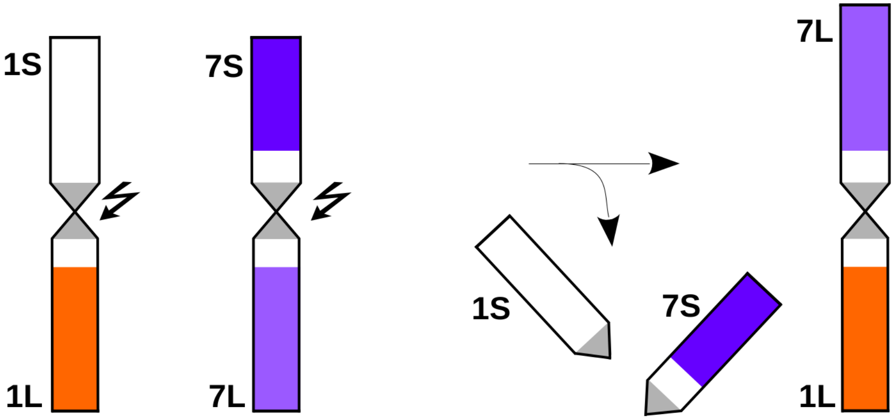

### Supplementary Figure S3

**Supplementary Figure S3.**

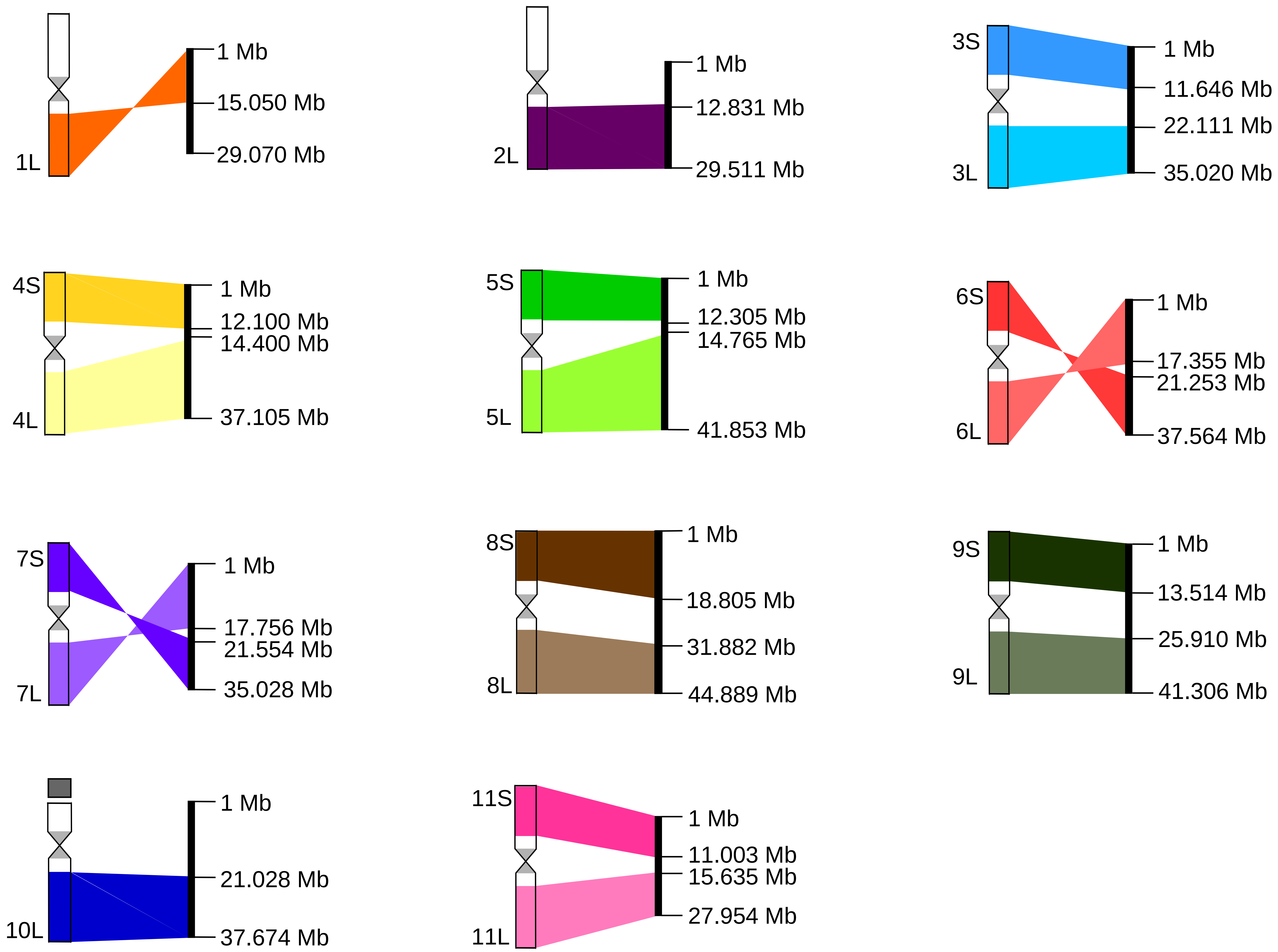
