## Supplementary Table S1 for "Chromosome painting in cultivated banana and their wild relatives (*Musa* spp.) reveals differences in chromosome structure"

| | Diploid accessions | | | | | | | | | | | | | | Triploid accessions | | | | | | | | |
|---|---|---|---|---|---|---|---|---|---|---|---|---|---|---|---|---|---|---|---|---|---|---|---|
| | M. acuminata ssp. (AA) | | | | | | | | Mchare cultivars (AA) | | | M. balbisiana (BB) | | M. schizocarpa (SS) | AAA | | | | AAB | | | ABB | |
| | | | | | | | | | | | | | | | Cavendish | Gros Michel | EAHB | | Plantains | | | Pelipita | Saba |
| Chromosome change | *malaccensis* **'Pahang'**** | *banksii* **'Bansksii'** | *microcarpa* **'Borneo'** | *Zebrina* **'Maia Oa'** | *burmannicoides* **'Calcutta 4'** | *burmannica* **'Tavoy'** | *siamea* **'Pa Rayong'** | ***P. lilin*** | Huti White | Huti (Shumba nyeelu) | Ndyali | Tani** | Pisang Klutuk Wulung | Schizocarpa** | Poyo | Gros Michel | Imbogo* | Kagera | 3 Hands Planty | Amou* | Obino l'Ewai | Pelipita | Saba sa Hapon |
| 3S/8L + 8S/3L | | | | •• | | • | | | • | • | • | | | | • | • | •• | •• | | | | | |
| 9S/1L + 1L/9S | | | | | •• | • | •• | | | | | | | | | | | | | | | | |
| 8L/2L | | | | | •• | • | •• | | | | | | | | | | | | | | | | |
| 7S/8S + 7L/8L | | | | | | • | | | | | | | | | | | | | | | | | |
| 3L/4S | | | | | | | • | | | | | | | | | | | | | | | | |
| 4L/1L + 1L/4L | | | | | | | | • | • | • | • | | | | • | • | | | | | | | |
| 7S/1L + telocentric 7L | | | | | | | | | | | | | | | • | • | | | | | | | |
| 1L/3L (B genome specific) | | | | | | | | | | | | •• | •• | | | | | | • | • | • | •• | •• |
| 7L/1L + missing 7S and 1S | | | | | | | | | | | | | | | | | • | | | | | | |
| missing chromosome 2 | | | | | | | | | | | | | | | | | | | | • | | | |

* aneuploid clones

** data from Šimoníková *et al*. (2019)
